# Paternal genomic resources from the YanHuang cohort suggested a Weakly-Differentiated Multi-source Admixture model for the formation of Han’s founding ancestral lineages

**DOI:** 10.1101/2023.11.08.566335

**Authors:** Zhiyong Wang, Mengge Wang, Kaijun Liu, Haibing Yuan, Shuhan Duan, Yunhui Liu, Lintao Luo, Xiucheng Jiang, Shijia Chen, Lanhai Wei, Renkuan Tang, Liping Hu, Jing Chen, Xiangping Li, Qingxin Yang, Yuntao Sun, Qiuxia Sun, Yuguo Huang, Haoran Su, Jie Zhong, Hongbing Yao, Libing Yun, Jianbo Li, Junbao Yang, Yan Cai, Hong Deng, Jiangwei Yan, Bofeng Zhu, 10K_CPGDP, Kun Zhou, Shengjie Nie, Chao Liu, Guanglin He

## Abstract

The large-scale human genome revolution and rapidly advanced statistical innovation have updated our understanding of the fine-scale and complex genetic structure, the entire landscape of genetic diversity and the evolutionary trajectories of spatiotemporally different ancients and ethnolinguistically diverse modern populations. Recent ancient DNA research provided a detailed and complex admixture picture of ancient Europeans but limited insights into East Asians as the few available genomes. Y-chromosome variations in the male-specific regions, served as molecular archaeological tool, have unique evolutionary features that can be utilized to reconstruct the origin and subsequent interaction of ancient East Asian paternal lineages. We launched the YanHuang cohort using our designed highest-resolution capture sequencing panel to explore the detailed evolutionary trajectory of the Han Chinese, one of the largest ethnic groups in the world. We reported one of the largest uniparental genomic resources and observed multiple founding paternal lineages dominant in ancient western Eurasian, Siberian and East Asian participating in the formation of the gene pool of the Han Chinese. We identified fine-scale paternal genetic structure correlated with different patterns of ancient population interaction and geographical mountain barriers (Qinling-Huaihe line and Nanling Mountains), suggesting isolation-enhanced and admixture-introduced genetic differentiation enhanced the complexity of the Han Chinese genomic diversity. We observed a strong direct correlation between the frequency of multiple founding lineages of the Han Chinese and the proportion of subsistence-related ancestry sources related to western pastoralists, Holocene Mongolian Plateau people and ancient East Asians, reflecting the ancient migration events contributed to our identified patterns of Chinese paternal genomic diversity. We finally provided one novel and the most plausible admixture-by-admixture model, the Weakly-Differentiated Multi-Source Admixture model, as the major genetic mechanism to illuminate our observed pattern of complex interactions of multiple ancestral sources and landscape of the Han Chinese paternal genetic diversity. Generally, we presented one large-scale uniparental genomic resource from the YanHuang cohort, portrayed one novel admixture formation model and presented the entire genomic landscape with multiple ancestral sources related to ancient herders, hunter-gatherers and farmers who participated in the ancestral formation of the Han Chinese.

## Introduction

East Asia, situated at the crossroad connecting America and the Pacific Islands, harbors a wealth of ethnolinguistic diversity and is an essential region for studying human evolution (Cavalli-Sforza 1998). Decades of genetic study have provided valuable insights into the source, formation and divergence of modern East Asians (Consortium et al. 2009; GenomeAsia 2019; Chen et al. 2022). The initial peopling history of East Asia occurred tens of thousands of years ago and unfolded along two distinct migration routes, the northern route and the southern route, contributing to the basal gene pool of the ancestor of East Asians (Jin and Su 2000; Zhong et al. 2011). The diverse patterns of genetic variations observed in present-day East Asians were shaped by different factors, such as population admixture, cultural and social practices, geographic isolation and others, indicating their different and complex population histories (Cao et al. 2020). Meanwhile, recent advances in ancient DNA investigation refreshed our understanding of the evolutionary history of ancient East Asians. Genetic continuity was evidenced via ancient DNA from the Amur River Basin and Tibetan Plateau and complex population interactions and admixture processes or key migration events were observed in core central regions of East Asia (Jeong et al. 2016; Mao et al. 2021; Wang et al. 2021a; Wang et al. 2021d; Liu et al. 2022a). Mao et al. identified Tianyuan/Yana-related Paleolithic lineages that contributed to the gene pool of Siberians and northern East Asians in the pre-last glacial maximum era and reported the genetic turnover between pre-Neolithic AR33K and AR19K and genetic continuity of Neolithic and modern Tungusic people (Mao et al. 2021). Ning et al. reported ancient DNA from northern China and identified the population genetic interaction associated with diverse subsistence strategies (Ning et al. 2020). Yang et al. reported genetic stratification between ancient northern East Asians (ANEA) and ancient southern East Asians (ASEA) since the early Neolithic period and Wang et al. identified the consistent southward gene flow from the Yellow River Basin (YRB) to Fujian and Guangxi Neolithic and historical populations (Yang et al. 2020; Wang et al. 2021d). McCol and Lipson et al. further identified southward gene flow from South China to Southeast Asia and even Oceania associated with spatiotemporally diverse ancient population expansion and migration (Lipson et al. 2018; McColl et al. 2018). Ancient western Eurasian Yamnaya and Afanasievo people also influenced the genomic diversity of ancient Eastern Eurasian Steppe (AEES) populations and Xinjiang people (Damgaard et al. 2018; Jeong et al. 2020; Zhang et al. 2021a; Kumar et al. 2022). Generally, complex interaction and admixture scenarios between spatiotemporally different ancient populations reshaped the genetic landscape of eastern Eurasians, consistent with the dispersal of ancient Chinese agriculture and pastoralist technological innovation and language families (Wang et al. 2021a). However, the extent to which these autosome-evidenced ancient population evolutionary events influenced the uniparental gene pool of modern East Asians and the fine-scale uniparental genetic history of Han Chinese populations remain unclear and need to be examined comprehensively.

The Han Chinese, one of the largest ethnic groups globally and spreading in geographically diverse regions of East Asia, have attracted geneticists to unravel their genetic diversity and population history over two decades (Wen et al. 2004a; Xu et al. 2009; Chiang et al. 2018; Cao et al. 2020; Cheng et al. 2023). From the perspective of genetics, archaeology and linguistics, the origin of Sino-Tibetan (ST) languages was associated with the Neolithic farmers who cultivated millet in the upper-middle YRB (Zhang et al. 2019; Wang et al. 2021a; Liu et al. 2022b) and the dispersal of ST languages during the Neolithic period aligned with the farming-and-language-dispersal hypothesis (Diamond and Bellwood 2003; Bellwood 2005). These early millet agriculturalists significantly contributed to the major genetic makeup of present-day Han Chinese populations (Wang et al. 2021a). Historically documented successive waves of dispersal of the Han Chinese and interactions with surrounding ethnolinguistically diverse populations indicated the complexity of the genetic history of the Han Chinese (Cavalli-Sforza et al. 1994; Wang 1994; Ge et al. 1997). Xu et al. provided anthropologic clues for the formation of the Han Chinese and proposed the “snowball theory” to illuminate the cultural formation of the Han Chinese, in which the Han Chinese rolled like a snowball, absorbing the cultural and genetic elements from surrounding ethnically different populations (Xu 1999; Xu 2012). Using the uniparental markers, Wen et al. revealed the southward demic diffusion of the Han Chinese with sex-biased admixture as the differentiation of the genetic contribution of northern Han to southern Han between the paternal and maternal lineages (Wen et al. 2004a). Subsequently, the phylogeographic pattern observed in the Y-chromosome presented that the genetic component of the Han Chinese was intricate and diverse, along with different paternal lineages (Xue et al. 2006; Xue et al. 2008; Zhong et al. 2011; Lang et al. 2019; Song et al. 2019; Tao et al. 2023). Meanwhile, autosomal DNA-based research identified different ancestral components among geographically diverse Han Chinese populations (Chen et al. 2019; He et al. 2020; He et al. 2021; Yao et al. 2021; He et al. 2022; Wang et al. 2022c). However, due to the restriction of sampling size and the paucity of ancient DNA data in the East Asian region, the genetic history and relationship between the Han Chinese and other ancient East Asian populations remain largely elusive. Similarly, the extent to which and how the past populations with different subsistence strategies, such as indigenous agriculture and neighboring nomadism, have contributed to the paternal genetic makeup of the Han Chinese remain uncertain since the Holocene.

Previous studies leveraged different molecular markers to explore the Han Chinese population structure, uncovering their distinct genetic relatedness and history patterns. Based on the early genome-wide array data, the researchers observed the North-to-South and East-to-West genetic divergences of the Han Chinese and identified its migration history followed the isolation-by-distance model that the genetic relatedness decreased as the geographic distance (Chen et al. 2009; Xu et al. 2009; Chiang et al. 2018). The high-throughput sequencing resources further advanced our understanding of the different population stratification among the Han Chinese, implying the diverse demographic history of geographically different East Asians (Liu et al. 2018; Cao et al. 2020; Zhang et al. 2021b; Cong et al. 2022; Qiu et al. 2022; Wang et al. 2022a; Cheng et al. 2023; Yu et al. 2023a). Unlike the nuclear genome data, the uniparentally-inherited markers can provide a unique evolutionary insight into the sex-specific genetic landscape and demographic history as their features of non-recombination and male/female-specific inheritance (Kutanan et al. 2019; Karmin et al. 2022; Li et al. 2023). Previous uniparental genetic evidence based on the limited number of markers identified the North-to-South cline of Han Chinese’s paternal and maternal genetic structure (Su et al. 1999; Yao et al. 2002; Wen et al. 2004a; Lang et al. 2019). Moreover, with the comprehensive sampling coverage, Li et al. unveiled the fine-scale matrilineal genetic divergence of the Han Chinese related to the river barriers and underscored the significant influence of agriculture technology innovation on shaping the matrilineal genetic variations (Li et al. 2019b). Although the prior research on the paternal genetic architecture of the Han Chinese had been substantially conducted (Wen et al. 2004a; Xue et al. 2008; Lang et al. 2019; He et al. 2023a), the restricted number of Y-SNPs and/or limitation of sampling bias have hindered our understanding of the picture of fine-scale paternal population structure and the critical factors in shaping the paternal genetic landscape as the patrilocality and matrilocality residence models possess differentiated role in the human Y-chromosome genetic diversity and variation spectrum.

Consequently, to better characterize the fine-scale paternal genetic structure and evolutionary history of the Han Chinese and explore their possible formation mechanism, we developed a high-resolution YHseqY3000 panel based on the uniparental genomic database and the highest-resolution phylogeny tree constructed in the 10K Chinese People Genomic Diversity Project (10K_CPGDP) (He et al. 2023b). We conducted the YanHuang cohort (YHC) and reported YHC uniparental genomic resources among 5,020 unrelated Han Chinese individuals from 29 administrative provinces of China using our developed YHseqY3000 panel (**Figure 1a**). We have presented, so far, one of the most extensive and comprehensive landscapes of Y-chromosomal genetic variations and the haplogroup frequency spectrum (HFS) in the Han Chinese. We explored the fine-scale paternal genetic structure of the Han Chinese as well as the possible influencing factors on the landscape of Han Chinese paternal genetic diversity, including cultural elements, river separation and mountain barriers. We also dissected the extent of diverse founding lineages introduced by extensive admixture events among previously isolated or differentiated sources that took part in the formation of the Han Chinese since the Holocene and proposed the **Weakly-Differentiated Multi-Source Admixture (WDMSA)** model that stated the multi-sources related to deep-in-time diverged but spatiotemporally and genetically weakly connected or recently admixed ancient western Eurasian herders, Siberian hunter-gatherers and East Asian millet and rice farmers participated in the formation of the Han Chinese. In aggregate, we provided a comprehensive insight into the genetic history of the Han Chinese, summarized and built one new genetic admixture-by-admixture model to illuminate the evolutionary process and underscored two important evolutionary forces, including isolation-enhanced and admixture-introduced genetic differentiation, in shaping the Han Chinese paternal genetic landscape.

**Figure 1.**
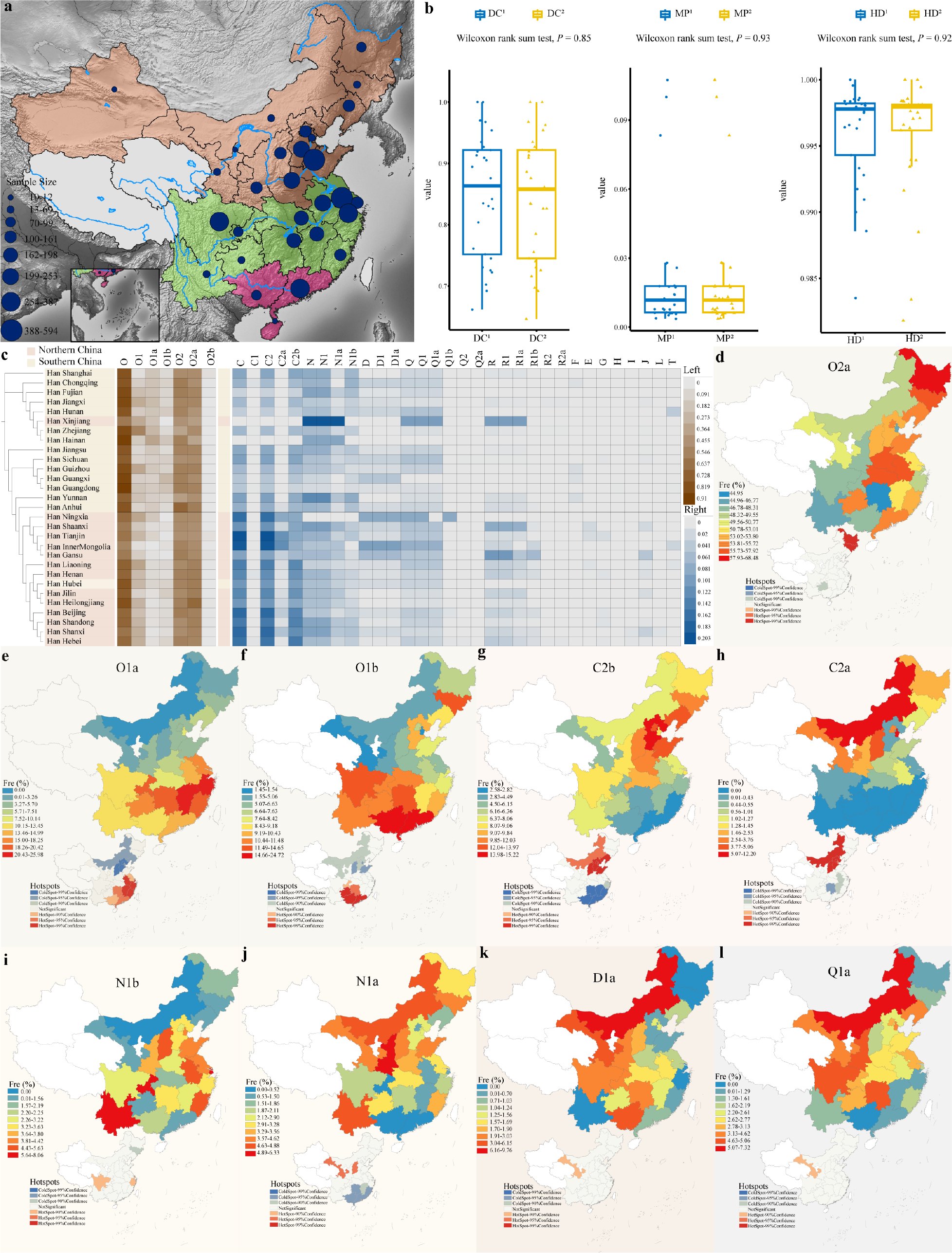
The geographical location of the newly collected samples and the haplogroup frequency distribution at the third level. **(a)** The geographical distribution of the YanHuang cohort samples from 29 administrative provinces in China. The position of the circle represents the sampling region and size indicates the sample size. The color of background indicated the different groupings according to the geographic barriers of Qinling-Huaihe line and Nanling Mountains. The white background presented the paucity of sampling. **(b)** The boxplots of DC (Discrimination capacity), MP (Match probability) and HD (Haplotype diversity) based on haplogroup and haplotype among the Han Chinese. The superscripts of 1 and 2 present haplotype and haplogroup, respectively. **(c)** The heat map of the haplogroup frequency spectrum within the third level among the Han Chinese. The topology of the tree is based on the F_ST_ calculated by the haplogroup frequency matrix at the third level. **(d-l)** The geographical distribution heatmap of the haplogroup O2a, O1a, O1b, C2b, C2a, N1b, N1a, D1a and Q1a at the third level among the Han Chinese; different colors represent the haplogroup frequency among the Han Chinese (top). The spatial autocorrelation analysis is based on the corresponding haplogroup frequency (bottom); HotSpot indicates the high-value spatial clustering, while ClodSpot indicates the low-value spatial clustering. Different colors present different confidence intervals. Due to the small total number of Xinjiang (10), Ningxia (12) and Hainan (11), these data were excluded.

## Results

### Paternal genetic diversity of the Han Chinese observed in the pilot work of the YanHuang cohort

We launched the YanHuang cohort to capture the entire landscape of paternal genetic diversity of geographically different Han Chinese populations and generated 5,020 targeted Y-chromosome sequences, including 3,002 phylogenetic informative SNPs (PISNPs) in our developed YHseqY3000 panel, from 29 different administrative provinces **(Figure 1a and Table S1)**. After the quality control, we observed 1,899 distinct haplotypes and 1,766 terminal haplogroups, which were allocated based on our developed forensic phylogenetic tree **(Table S2)**. The haplotype/haplogroup-based parameters within the Han Chinese, such as haplotype/haplogroup diversity ranging from 0.9835 to 1.0000 and from 0.9818 to 1.0000, respectively, showed similar and strong values **(Figure 1b and Table S3)**, indicating that the YHseqY3000 panel had a rather powerful performance in haplogroup classification and variation capture.

To investigate the frequency distribution of founding lineages or dominant haplogroups consistent with historical coherence of haplogroup Nomenclature, we used HaploGrouper to classify haplogroup based on the International Society of Genetic Genealogy (ISOGG) Y-DNA Haplogroup Tree 2019-2020 (version 15.73) and finally identified 545 definitive haplogroups among 5,020 unrelated samples **(Table S2)**. We first assessed the geographical distribution of major haplogroups at the third level and generated the HFS of the Han Chinese **(Figure 1c)**. Haplogroup O2a (52.85%) contributed significantly to the paternal gene pool of the Han Chinese, followed by O1a (12.47%), O1b (9.86%), C2b (8.71%), N1b (3.57%), N1a (2.77%), Q1a (2.77%), D1a (1.43%) and C2a (1.08%) **(Table S4)**. To explore the distribution patterns of these haplogroups in detail, we generated the geographical distribution heatmap using ArcGIS 10.8 **(Figures 1d-l)**. The dominant haplogroup O2a had the highest frequency in northern and northeastern China **(Figure 1d)**, including Heilongjiang (68.47%), Hubei (57.92%) and Anhui (56.78%) **(Table S4)**. The majority of its sub-lineages consisted of O2a2b1a1-M117 (16.91%), O2a2b1a2a1a-F46 (9.92%) and O2a1b1a1a1a-F11 (10.66%) **(Table S4)**. O1a-M119 was distributed mainly in southern and southeastern China **(Figure 1e)**, including Zhejiang (25.98%), Jiangxi (23.47%) and Fujian (20.42%) **(Table S4)**. In particular, the sub-lineage of O1a-M119 was mainly composed of O1a1a1a1a1a1-F492 (5.40%) **(Table S4)**. O1b-M268 was mainly distributed in southern China **(Figure 1f)**, especially in Guangxi (24.72%), Guangdong (19.89%) and Hunan (14.65%) **(Table S4)**. Interestingly, the distribution of the sister sub-haplogroups, O1b1a1-PK4 and O1b1a2-Page59, were correlated with geographical distances and showed a latitude-dependent gradient, with O1b1a1-PK4 more in southern China and O1b1a2-Page59 more in northern China **(Figure S1)**. C2b was mainly found in northern China **(Figure 1g)**, including Hebei (15.21%), Shandong (13.97%) and Beijing (13.53%) **(Table S4)**. We also noticed that C2a had a high frequency in northern China **(Figure 1h)**, such as Inner Mongolia (12.20%) and Tianjin (11.59%) and was distributed rarely in southern China compared to C2b **(Table S4)**. N1b was mainly distributed in southern and southwestern China **(Figure 1i)**, including Yunnan (8.06%), Chongqing (7.07%) and Shanghai (6.57%) **(Table S4)**. N1a was mainly found in northern China **(Figure 1j)**, such as Shaanxi (6.32%) and Inner Mongolia (4.88%) **(Table S4)**. Notably, N1a occupied a certain proportion in Yunnan (4.84%) **(Table S4)**. D1a had a high frequency in the surrounding areas of the Tibetan Plateau, such as Inner Mongolia (9.76%) and Gansu (6.15%) **(Figure 1k)**. D1a was prevalent among Tibetans and the dominant paternal lineage of the Tibeto-Burman-speaking (TB) populations (Shi et al. 2008; Qi et al. 2013), suggesting interaction between the Han Chinese and TB populations. Q1a was mainly distributed in northern China **(Figure 1l)**, including Inner Mongolia (7.32%) and Shaanxi (5.06%) **(Table S4)**. Most of its sub-lineages were Q1a1a-M120 (2.75%) **(Table S4)**. The rare deep-rooted haplogroup F (0.08%) was entirely distributed in southwestern China **(Table S4)**, consistent with the southern route of the prehistorically northward migration of modern humans in East Asia (Su et al. 1999). Other non-Chinese-specific haplogroups, such as R, E, G, H, I, J and L, were mainly observed in northern and northwestern China **(Table S4)**, providing clues from the Holocene trans-Eurasian connection. These identified West Eurasian and Central/South Asian-specific haplogroups directly reflected the east-west gene flow within the Eurasian continent along prehistoric trans-Eurasian communication and historic migration along the ancient Silk Road (Karafet et al. 2008; Zhong et al. 2011). In aggressive, the paternal lineages of the Han Chinese were diverse and multivariate and these results also represented the geographically specific distribution of haplogroups within the Han Chinese.

### Isolation-enhanced genetic differentiation influenced the paternal genetic architecture of the Han Chinese

To explore genetic relatedness within the Han Chinese, we performed principal component analysis (PCA) on 4,987 individuals from 26 of the 29 sampling administrative provinces of China based on the haplogroup frequency at the fourth level **(Figure 2a-c)**. When we grouped populations by geography, the PC1 separated the Han Chinese into northern and southern Han Chinese, with the Qinling-Huaihe line as the geographical boundary **(Figure 2a)**. We also found that PC1 was highly correlated with latitude (*R* = - 0.82, *P* < 0.001) while not correlated with longitude **(Figure 2d)**. Of note, the longitude showed a significant correlation with O2a (*R* = 0.56, *P* < 0.001), which might result from the recent westward migration of the Han Chinese since 1949 (Liang and White 1996) **(Figure 2d)**. In addition, the correlation analysis between PC1 and frequency of O1a (*R* = 0.96, *P* < 0.001) and C2b (*R* = - 0.69, *P* < 0.001) was strongly statistically significant **(Figure 2d)**. It appeared that O1a had a high frequency in the southern populations clustering on the right of the PC map and decreased to a low frequency in the northern populations on the left, whereas the frequency of C2b showed a contrasting pattern. The North-South stratification was also revealed by the neighbor-joining tree and the genetic distance matrix based on pairwise genetic distances (F_ST_) as well as the correlation analysis between the pairwise genetic distances among 26 Han Chinese populations **(Figures 2d-f)**.

**Figure 2.**
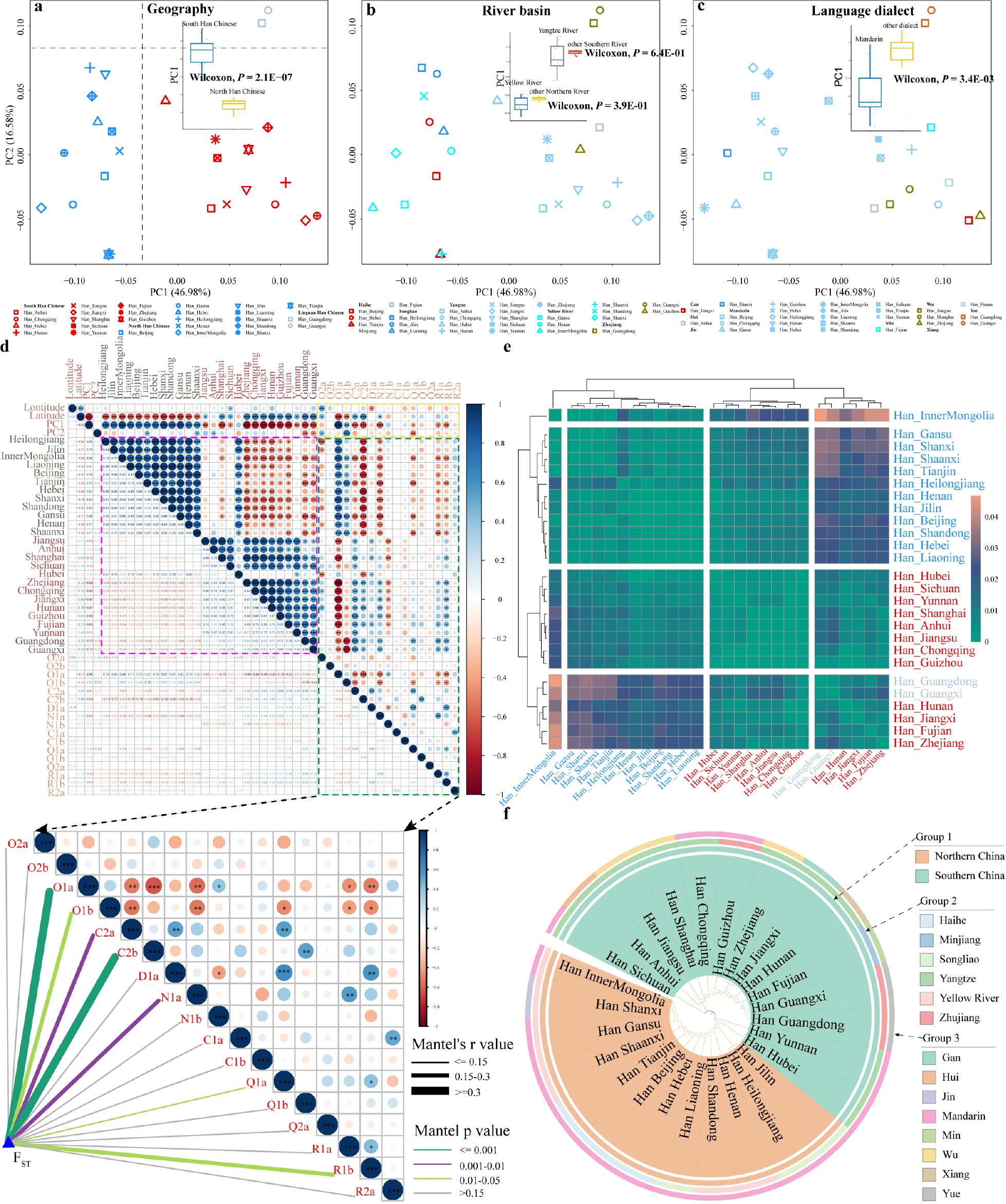
Paternal genetic structure and correlation analysis. **(a-c)** PCA of the Han Chinese based on the haplogroup frequency matrix at the fourth level, excluding the Xinjiang, Ningxia and Hainan due to potential sampling bias, same to following pictures. Populations are grouped by (a) geographical boundaries of the Qinling-Huaihe line and Nanling Mountains, (b) different river valleys and (c) different language dialects. **(d)** Correlation analysis is shown between longitude, latitude, PC1, PC2, pairwise genetic distance and haplogroup frequency among the Han Chinese. The pairwise genetic distance was related to each haplogroup composition by the Mantel test (bottom). Edge color indicates the statistical significance and Edge width presents the Mantel’s r statistic. **(e)** The heat map of the matrix based on the pairwise genetic distance among the Han Chinese. **(f)** The neighbor-joining tree based on the pairwise genetic distance among the Han Chinese. The different colors present different groupings.

Southern Han Chinese exhibited a distinct substructure in the PCA clustering pattern, in which PC2 effectively separated Guangdong and Guangxi populations from other southern Han Chinese, with the Nanling Mountains as another geographical boundary **(Figure 2a)**. Hence, we categorized Guangdong and Guangxi individuals as Lingnan Han in the subsequent genetic analysis. Notably, the pairwise genetic distances observed between the merged northern and southern Han without Lingnan Han (0.0102) were equal to the estimated values between the merged southern Han and Lingnan Han (0.0100), further conforming Lingnan Han to another genetic sub-clade **(Table S5)**. PC2 coordinates were not correlated with longitude or latitude but possessed a strong correlation with the frequency of O1b (R = 0.74, *P* < 0.001) **(Figure 2d)**. In the neighbor-joining tree, we observed that the Central Han (Anhui, Jiangsu and Shanghai) were clustering together **(Figure 2f)**. However, the Central Han people were not clearly separated from other southern Han **(Figure 2a)**. Our observed genetic similarity and difference patterns were further confirmed via the multidimensional scaling (MDS) analysis **(Figure S2)**. In addition, we did not identify isolation by distance pattern in the paternal genetic structure (r = 0.0006, *P* = 0.462), which was evidenced in the autosomal DNA using the Mantel test (Chiang et al. 2018). The differences in the genetic features inferred from the autosomal and uniparental genetic legacy showed different genetic influences of patrilocality and matrilocality residence models on the gene pool of the Han Chinese. Additionally, we explored the genetic affinities among the Han Chinese separated along different river valleys and language dialects **(Figures 2b and c)**. Populations from the northern River (YRB and other Northern Rivers) exhibited significant differentiation from others in the southern River (Yangtze River and other Southern Rivers) along PC1 **(Figure 2b)**. Notably, Mandarin-speaking populations distinguished substantially from others along PC1 (Wilcoxon rank sum test, *P* = 3.40E-3) **(Figure 2c)**, underscoring the association between cultural dialects and Y-chromosome founding lineages. In summary, according to the results above, we identified three population substructures based on the large-scale Y-chromosome sequences, including North Han, South Han, and Lingnan Han **(Figures 2a and S2)**.

To further dissect the genetic differentiation and the factors that influenced the paternal genetic structure of the Han Chinese, we performed the analysis of molecular variance (AMOVA) by classifying the Han Chinese into different catalogs based on the different geographical regions, river valleys, and language dialects **(Table 1)**. All genetic variances from the within-population level accounted for 1.02% (*P* < 0.001) **(Table 1)**. When we classified the Han Chinese into northern and southern Han, the northern Han (0.18%, *P* > 0.001) showed greater genetic homogeneity than the southern Han (0.67%, *P* < 0.001) **(Table 1)**, which indicated the different patterns of genetic differentiation between geographically diverse regions. The significant among-group variation (1.14%, *P* < 0.001) was presented in the grouping that categorized the Han Chinese into North Han, South Han, and Lingnan Han, which was higher than other groups, such as groups categorized by the four/seven geographical regions, river valleys and language dialects **(Table 1)**. Interestingly, the among-group variation of the grouping cataloged by three river valleys showed a relatively high value (1.09%, *P* < 0.001) when compared with the grouping of northern Han and southern Han (1.06%, *P* < 0.001) **(Table 1)**. These results showed that isolation-enhanced genetic differentiation presented by the geography boundaries (Qinling-Huaihe line and Nanling Mountains) and the river valleys (Yellow River, Yangtze River, and Zhujiang River) played a pivotal role in shaping the paternal population structure of the Han Chinese.

**Table 1.** Results of AMOVA analysis.

| Groupings | Number of Populations | Number of Groups | Percentage of Variations (%) |  |  |
| --- | --- | --- | --- | --- | --- |
|  |  |  | Among-Groups | Among Populations within Groups | Within-Populations |
| Total | 26 | 1 | - | 1.02** | 98.98** |
| Southern China vs. Northern China<sup>a</sup> | 26 | 2 | 1.06** | 0.47** | 98.46** |
| Southern China vs. Northern China versus Southwestern China vs. Northwestern China<sup>b</sup> | 26 | 4 | 0.88** | 0.44** | 98.68** |
| Central vs. eastern vs. Northeastern vs. Northern vs. Northwestern vs. Southern vs. Southwestern<sup>c</sup> | 26 | 7 | 0.31 | 0.71** | 98.98** |
| South Han vs. North Han vs. Lingnan Han<sup>d</sup> | 26 | 3 | 1.14** | 0.32** | 98.54** |
| Yangtze River vs. Yellow River vs. Zhujiang River vs. Haihe River vs. Songliao River vs. Minjiang River<sup>e</sup> | 26 | 6 | 0.96** | 0.31** | 98.73** |
| Yangtze River vs. Yellow River vs. Zhujiang River<sup>f</sup> | 26 | 3 | 1.09** | 0.32** | 98.59** |
| Language Dialects<sup>g</sup> | 26 | 8 | 0.93** | 0.36** | 98.71** |
| Northern Han Chinese<sup>a</sup> | 14 | 1 | - | 0.67* | 99.33* |
| Southern Han Chinese<sup>a</sup> | 12 | 1 | - | 0.18** | 99.82** |
| Lingnan Han Chinese<sup>d</sup> | 2 | 1 | - | 0.17 | 99.83 |
| Central Han Chinese | 3 | 1 | - | -0.12 | 100.12 |

**Note:** The superior characters indicate different groupings. Detailed information about groupings is shown in Table S6. \*\*indicates $P < 0.01$ \*indicates $P < 0.05$ .

### The genetic connection and expansion events among founding populations with the different paternal lineages

To comprehensively investigate the genetic connection among different paternal lineages and between observed population substructures, we performed median-joining network analysis and reconstructed a parsimony phylogenetic tree using 5,020 Y-chromosome sequences **(Figures 3 and S3)**. We observed that nearly every central node and branch was contributed by geographically diverse populations, and the paternal lineages consisting of O2a, O1a, O1b, and C2b contributed to most of the paternal genetic framework **(Figures 3a and S3)**. Notably, the O1a node and branch predominantly comprised individuals from South Han, while Lingnan Han primarily represented the O1b node and branch. In contrast, the C2b node and branch primarily consisted of North Han **(Figures 3a and S3)**. We then generated the network focusing exclusively on O2a, O1a, O1b, C2b, C2a, N1a, N1b, Q1a, and D1a to explore the fine-scale genetic structure of these lineages comprehensively **(Figures 3b-c and S4-6)**.

**Figure 3.**
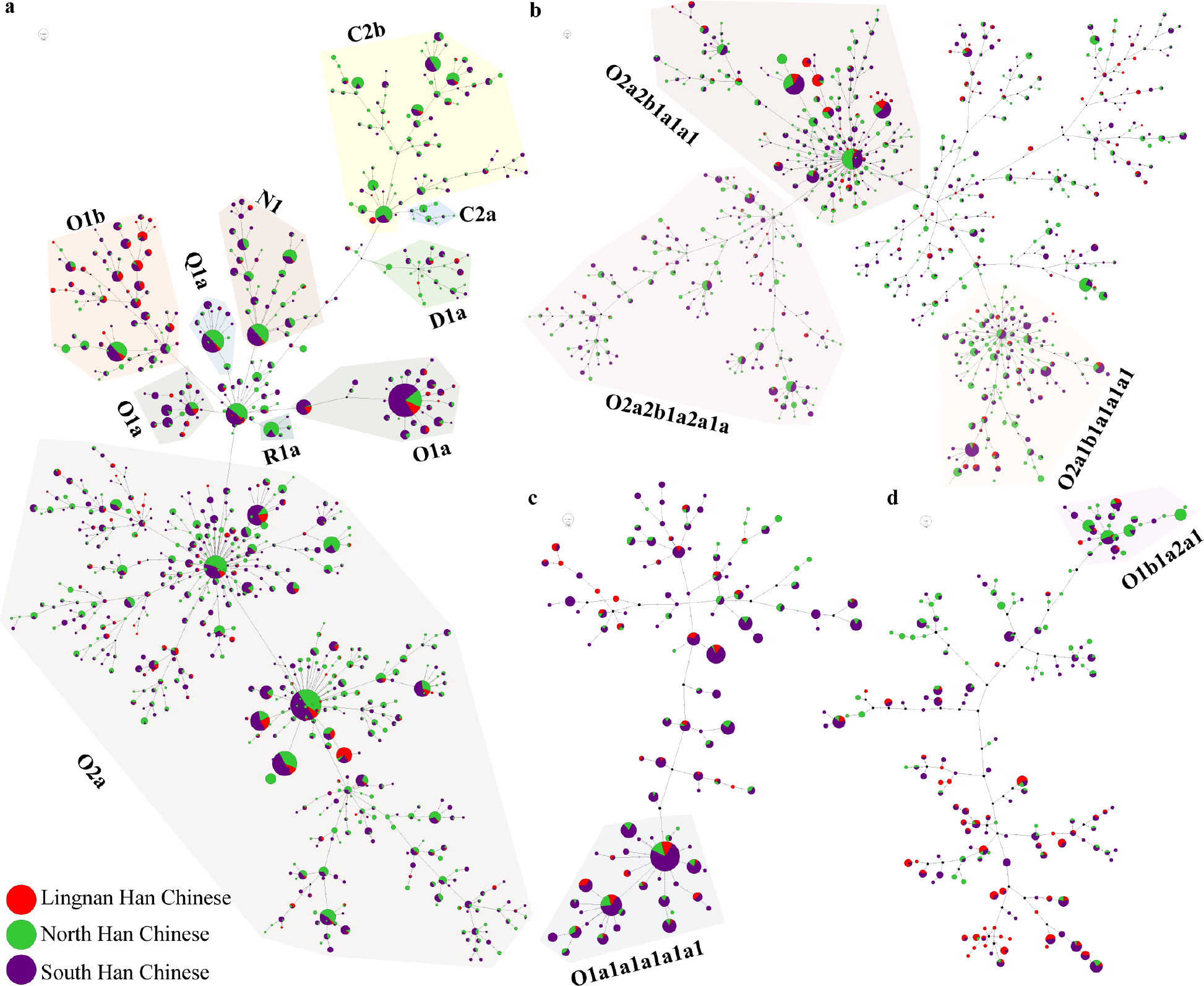
Median-joining network among the Han Chinese. **(a)** Median-joining network among all Han Chinese in our study. **(b)** Median-joining network among the Han Chinese attributed to O2a and its sub-lineages. **(c)** Median-joining network among the Han Chinese allocated to O1a and its sub-lineages. **(d)** Median-joining network among the Han Chinese classified to O1b and its sub-lineages. Different colors in the node present different groups. Different color backgrounds indicate different haplogroups and their sub-haplogroups.

Our results presented the star-like network of O2a2b1a1a1 (downstream of O-M117) and O2a1b1a1a1a1 (downstream of O-F11) **(Figure 3b)**, indicating the expansion of these lineages in the Han Chinese. However, we had not observed an apparent hierarchical expansion structure within O2a2b1a2a1a-F46, and the origin of the cluster was not located in the center **(Figure 3b)**, which might have indicated the expansion of O2a2b1a2a1a-F46 happened immediately after its advent. Our results reconciled the phylogenetic bifurcations of three Neolithic Super-Grandfathers of the Han Chinese (Yan et al. 2014), revealing deeper branches’ expansion in the Han Chinese. Ancient DNA analysis identified that O1a was first found in the 5,300-4,100-year-old Liangzhu ancient people in the surrounding region of the Yangtze River delta (Li et al. 2007). Previous studies found that O1a sub-lineages underwent a complex demographic history over the past 10,000 years, in which the Neolithic communities in Southeast China with O1a contributed to the gene pool of Sinitic/Tai-Kadai/Austronesian-speaking populations (Sun et al. 2021). The striking star-like network of O1a1a1a1a1a1-F492 implied that the haplogroup had undergone expansion in the Han Chinese, which was in line with the phylogenetic radiation in previous studies (Sun et al. 2021).

The O1b1a2-Page57 showed a high frequency in northeast China **(Figure S1)**, and its sub-lineage O1b1a2a1-F1759 presented a star-like network in the Han Chinese **(Figure 3d)**. Meanwhile, the previous population investigation found that O1b1a2-Page57 was mainly spread among Sinitic-speaking populations while absent in other ethnolinguistic populations (Lang et al. 2019; Song et al. 2019; Xie et al. 2019; Song et al. 2021; Wang et al. 2021b; Wang et al. 2021c; Wang et al. 2022b), and O1b1a2 was detected in the Neolithic people from Wanggou site (5,500-5,000BP) belonging to Yangshao culture (Ning et al. 2020), which might take part in the formation of the Han Chinese and finally became one of their founding paternal lineages. In addition, we also observed that the sub-lineages of Q1a and N1a showed a similar pattern **(Figures S5b and S6a)**, which was in keeping with the phylogenetic expansion in previous studies (Sun et al. 2019; Yu et al. 2023b). This evidence revealed the internal genetic connection and the expansion events within the Han Chinese. Meanwhile, these expansion events, which were also confirmed by previous phylogenetic studies, demonstrated that our designed panel can comprehensively capture the genetic diversity of Han Chinese populations.

### The effect of the admixture-introduced genetic differentiation on the formation of the Han Chinese

The observed paternal substructure of the Han Chinese defined the level of genetic differentiation and extent of the Y-chromosome variations, which provided clues to explore the differences in the composition of founding lineages. We first performed Fisher’s exact test to investigate which haplogroup showed a significant North-South frequency difference and observed that O1a, O1b, C2b, C2a, R1a, R1b, N1a, and J2a presented significant differences of haplogroup frequency distribution between southern and northern Han populations **(Table S7)**. We further performed the mantel test to explore the correlation between the pairwise genetic distances and haplogroup composition **(Figure 2d)**. Overall, O1a, O1b, C2b, C2a, N1a, and R1b significantly contributed to the discrepancy of the pairwise genetic distances between the Han Chinese populations **(Figure 2d)**. Analysis of spatial autocorrelation can investigate the pattern of regional clustering among these major haplogroups, which might relate to the corresponding haplogroup’s diffusion center **(Figures 1d-i)**. Results revealed a pronounced clustering pattern for O1a and O1b within southern China, where O1a clustered near the Yangtze River and O1b clustered in the Lingnan region. **(Figures 1e and f)**. A significant pattern of regional clustering in northern China was prominently observed for C2b and C2a **(Figures 1g and h)**. Nonetheless, when considering the prevalent haplogroup O2a and its major sub-haplogroup O-M117, O-F46, and O-F11 among the Han Chinese, the obvious regional clustering pattern above was not observed, which might indicate the extensive admixture of the three Neolithic Super-Grandfathers **(Figures 1e and S1)**. In general, taking into account the prevalence of these haplogroups within the Han Chinese, various statistical analyses had elucidated that the predominant influence on the observed differentiation within the paternal genetic structure of the Han Chinese was primarily attributed to the presence of O1a, O1b, and C2b haplogroups. Meanwhile, the genetic components of C2a, N1a, R1a, R1b, and J2a have also contributed to genetic differentiation within the Han Chinese.

The composition of the paternal lineages observed in the Han Chinese was diverse, and some of the ancient indigenous and incoming paternal lineages more or less contributed to their genetic differentiation. To distinguish the extent to which ancient populations in Eurasia contributed to the paternal genetic makeup of the Han Chinese since the Holocene, we performed the model-based ADMIXTURE analysis of 32 geographically diverse Han Chinese populations and ancient Eurasian populations to explore their basic admixture genetic profile **(Figure 4a)**. We found that the four-source admixture model best explained the genetic ancestry composition of the Han Chinese, including YRB-related ANEA ancestry (Pingliangtai_LN; green), ASEA-related ancestry (GaoHuaHua; orange), AEES-related ancestry (Mongolia_N_North; purple) and ancient Western Eurasian Steppe-related ancestry (AWES; Afanasievo; blue) **(Figure 4a)**. The ancestral component exhibited variations across geographically diverse Han Chinese populations **(Figure 4b)**. In short, the proportion of AEES/AWES/ANEA-related ancestry decreased from north to south, whereas the tendency of ASEA-related ancestry presented a contrasting pattern among the Han Chinese **(Figure 4b)**.

**Figure 4.**
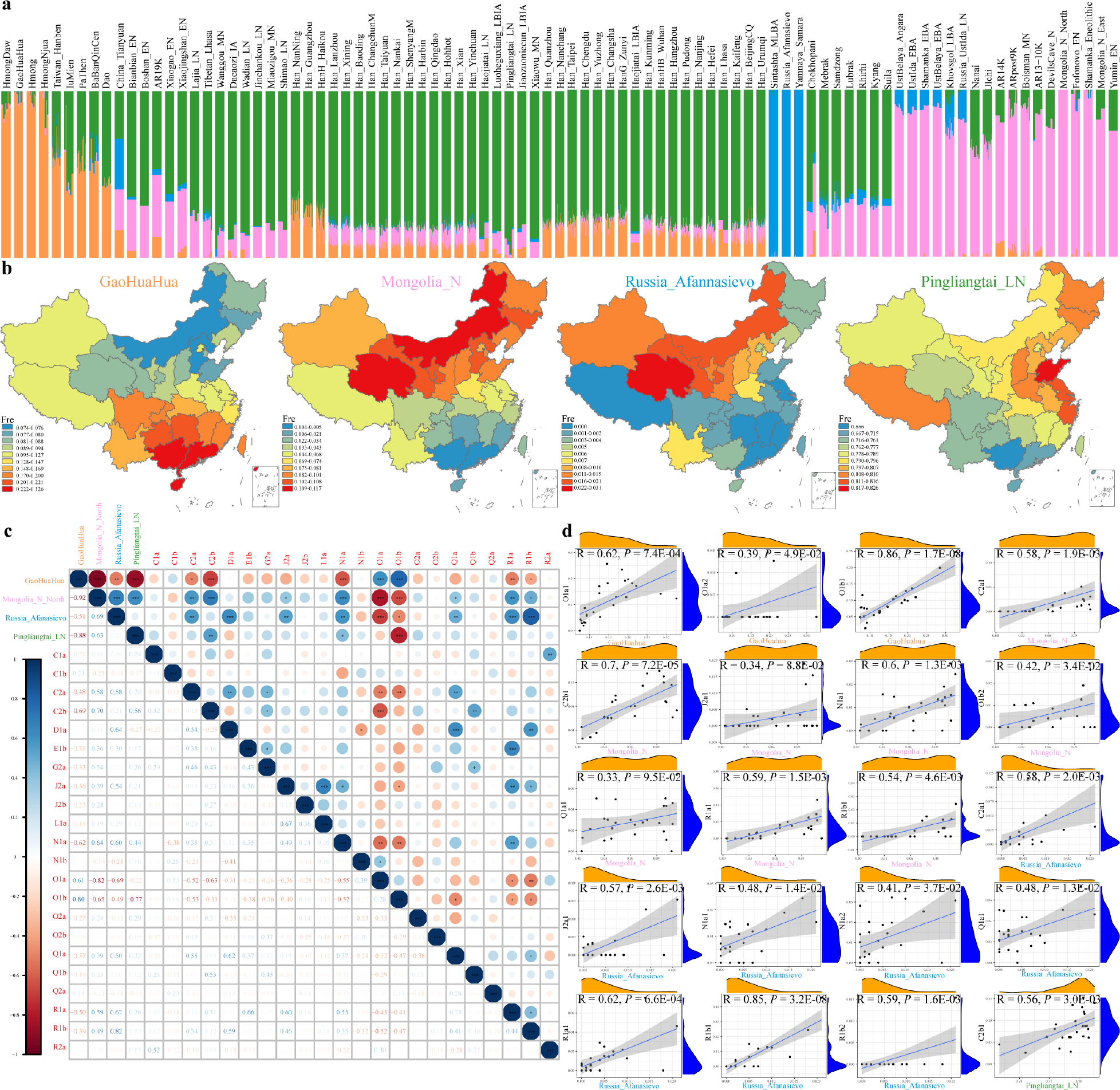
Correlation analysis between Admixture ancestry components and paternal lineages among the Han Chinese. **(a)** ADMIXTURE results presented the ancestry component of the Han Chinese with West Eurasia, Siberia and East Asia ancient individuals. **(b)** The distribution of the Admixture ancestry components among the Han Chinese. **(c-d)** The correlation analysis between the Admixture ancestry component and paternal lineages at the third and fourth level, respectively.

Previous genomic findings had evidenced the male-dominant sex-biased admixture in TB and Hui populations (Wen et al. 2004b; Ma et al. 2021). Consequently, if the proportion of estimated ancestral component demonstrates a significant positive correlation with the frequency of paternal lineages, it is plausible to consider that the expansion and admixture events related to the ADMIXTURE-inferred ancestral sources might have contributed to the incorporation of the paternal lineage into the formation of the targeted Han Chinese. Subsequently, we conducted a correlation analysis between the ADMIXTURE-inferred ancestry proportion and paternal lineage frequency in geographically different Han Chinese. We observed that ASEA-related ancestry composition was significantly related with O1a and O1b, AEES-related ancestry composition was correlated with C2a, C2b, J2a, N1a, Q1a, R1a and R1b, AWES-related ancestry composition was associated with C2a, D1a, J2a, N1a, Q1a, R1a and R1b, and ANEA-related ancestry composition was correlated with C2b and N1a **(Figure 4c)**. We further explored the correlation between the sub-lineages at the fourth level and genetic ancestry composition **(Figure 4d)**. The downstream of these lineages was also generally associated with corresponding genetic ancestry composition **(Figure 4d)**. Interestingly, we observed that these paternal lineages introduced by the corresponding ancestry-related populations largely contributed to the genetic differentiation of the Han Chinese **(Figure 2d and Table S7)**. In general, these admixture-introduced external paternal lineages contributed to genetic divergence among the northern and southern Han Chinese, confirming that the evolutionary force of population admixture within Eurasia played a vital role in the formation of the paternal genetic landscape of the Han Chinese.

### The weakly-differentiated multi-source admixture model for the origin of the Han Chinese

Recent genomic evidence has illuminated three ancestral sources related to local hunter-gatherers, western Eurasian Yamnaya pastoralists and Near East farmers who formed modern European people (Lazaridis et al. 2014). Ancient and modern genetic admixture modeling also illuminated that modern Indian people carried two primary ancestral sources of ancient northern Indian and ancient southern Indian related to the ancient pastoralist expansion and admixture with local residents (Reich et al. 2009). Admixture coalescent modeling of modern African also revealed that the weakly structured stems model contributed to the modern African origins (Ragsdale et al. 2023). Our genetic studies revealed that multiple sources contributed to the formation of the genetic diversity of the Han Chinese based on both autosomal ancestral components and the paternal lineages **(Figures 1c and 4a)**. From the ancient genetic landscape of paternal lineages, we could discover that some of the above four ancestry-related populations have shared paternal lineages since the Holocene. For example, the typical haplogroup R1a and R1b derived from AWES-related populations emerged in the AEES-related populations (Narasimhan et al. 2019; Wang et al. 2021a), such as Mongolia_LBA_MongunTaiga_3 (R1a) and Mongolia_EBA_Chemurchek (R1b). The N1a also emerged in the AWES-related populations (Mereke_MBA) and AEES-related populations (Munkhkhairkhan_MBA) (Narasimhan et al. 2019; Wang et al. 2021a). The ANEA-related lineage O2a appeared in the individuals (Mongolia_LBA_CenterWest_5) from the AEES-related populations (Ning et al. 2020; Wang et al. 2021a). The C2b was detected in the AEES-related populations (Russia_Siberia_Lena_EN) and ANEA-related populations (Shimao_LN and Miaozigou_MN) (Ning et al. 2020; Yu et al. 2020). Besides, we could observe that diverse ancestral sources spread identical paternal lineages into the gene pool of the Han Chinese when we analyzed the correlation between the lineage frequency and ADMIXTURE-based ancestry proportion. R1a, R1b and N1a correlated with AWES/AEES-related ancestry component, and C2b correlated with AEEA/ANEA-related ancestry component **(Figure 4c)**, which was consistent with present findings from the archaeogenetics among the Eurasia. The shared and correlated paternal lineages might indicate the potential gene flow between these ancestral sources.

Except for the observed shared paternal lineages between the aforementioned ancestry-related sources, we also tried to fit similar admixture patterns based on the autosomal genomic variations to illustrate the in-depth ancient divergence and recent connection of these ancestral sources in the Han Chinese formation processes. Consequently, we used ADMIXTOOLS2 analysis to explore further population divergence and gene flow between the Han Chinese and other ancient populations and to dissect the putative admixture model that explained the formation of the Han Chinese well **(Figure S7)**. The best model fitting of the admixture graph presented that the formation of the Han Chinese was fitted with 51% ancestry related to GaoHuaHua and 49% ancestry related to YRB millet farmers. We can also observe the ancient gene flow between Afanasievo herders and Mongolia plateau hunter-gatherers, between Mongolia_North_N and China_YR_LN from the admixture graph **(Figure S7)**, suggesting the early connection of these putative ancestral sources. Generally, our findings suggested that ancient East Asians, including ANEA and ASEA, ancient Siberians and ancient western Eurasian pastoralists with differentiated but connected genetic lineages, participated in forming modern Han Chinese (**Figure 5**). The native ancestry sources, represented by ANEA and ASEA-related populations, were major spread the O2 and O1 haplogroups and their sublineages, which had historical connections to the founding lineages of cultivation millet and rice farmers. Siberian hunter-gatherers with AEES-related ancestry components primarily disseminated the C and their sublineages through complex interaction and admixture with AWES-related populations and ANEA-related millet farmers. AWES-related pastoralists mainly brought R and their subhaplogroups into northwestern Han Chinese with the pastoralist expansion or Holocene trans-Eurasian cultural exchange and population admixture, which also exhibited a connection with AEES-related populations. We summarized the weakly differentiated but multiple connected ancestral sources presented by native and immigrant ancestry populations flowing extensive expansion and admixture processes as the WDMSA model (**Figure 5**). Our model suggested that the complex, multiple and multi-layered ancestral sources participated in forming geographically diverse modern Han Chinese populations. Our identified fine-scale paternal genetic structure also suggested that both isolation-enhanced and admixture-introduced genetic differentiation contributed to the genetic differences among North, South and Lingnan Han Chinese populations.

**Figure 5.**
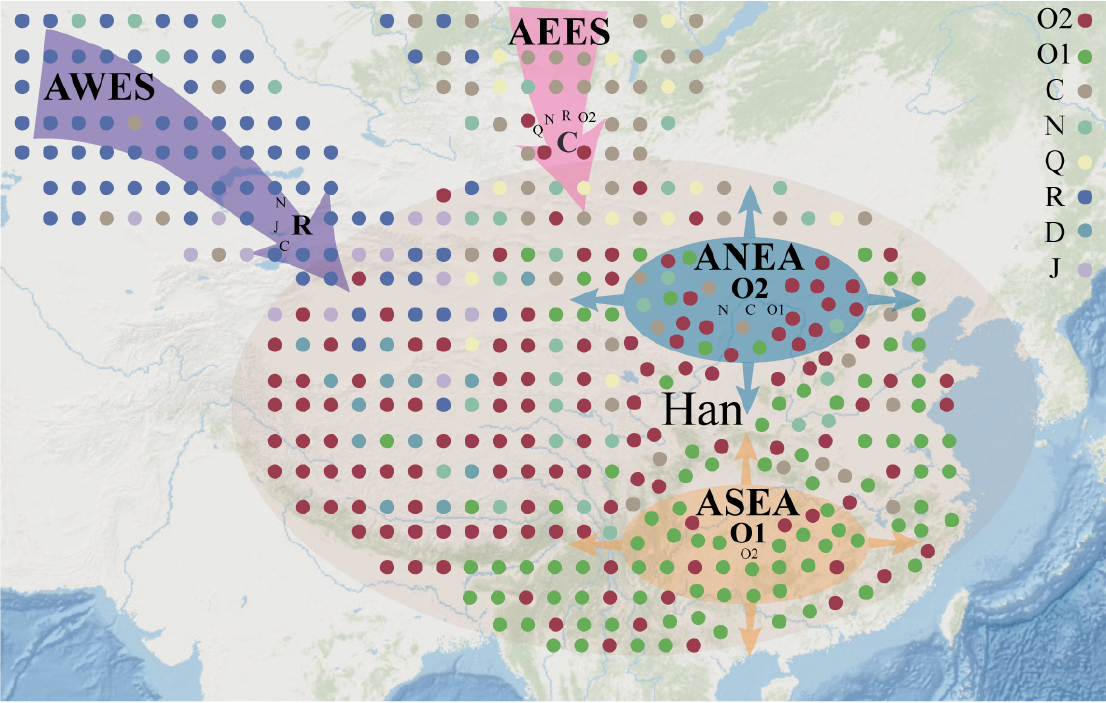
The simplified evolution pattern of the Han Chinese. The weakly-differentiated multi-source admixture model was proposed to illuminate the formation of the Han Chinese. The different colors of the circle indicate the different haplogroups. The abbreviations are presented as follows. **AEES**: Ancient Eastern Eurasia Steppe. **AWES**: Ancient Western Eurasia Steppe. **ANEA**: Ancient northern East Asians. **ASEA**: Ancient southern East Asians.

### The genetic relatedness between the Han Chinese and other ethnolinguistic populations in East Asia

To explore the genetic affinity and admixture signature between the Han Chinese and other ethnolinguistic populations in East Asia, we aggregated our data with the published available haplogroup information, resulting in a comprehensive dataset encompassing 10,481 individuals from 54 ethnolinguistically diverse populations belonging to four language families (Lang et al. 2019; Song et al. 2019; Xie et al. 2019; Wang et al. 2021b; Wang et al. 2021c; Song et al. 2022; Wang et al. 2022b; He et al. 2023a). We performed PCA based on haplogroup frequency at the fourth level, aiming to elucidate the genetic relationship among the combined populations. We observed that northern and southern Chinese populations were generally separated along the PC2 axis. Populations from northern China were predominantly positioned toward the lower part of the PC map, while populations from southern China were predominantly positioned toward the upper part **(Figure 6a)**. The particular distribution pattern underscored the genetic divergence between northern and southern Chinese populations, which was also conspicuously proved by the neighbor-joining tree and the distance matrix **(Figures 6b-d)**. Interestingly, we observed that the pattern of population clustering was generally relevant to linguistic affiliation **(Figures 6b-d)**, implying the association between the language family and paternal lineage. Furthermore, the neighbor-joining tree and the distance matrix showed that the Altaic-speaking populations from northern China were relatively close to the northern Han Chinese. In contrast, the TB/Tai-Kadai-speaking populations from southern China exhibited relative proximity to the southern Han Chinese **(Figures 6b-d)**, indicating the gene flow between the Han Chinese and surrounding ethnolinguistically diverse populations. In particular, Hui populations were relatively close to the Han Chinese **(Figure 6a)**, which was consistent with a previous study (Ma et al. 2021). We also noticed that the D1a has a high frequency in Tibetans, which was consistent with previous studies (Qi et al. 2013). R1a, R1b and R2a were mainly located in northwestern China, especially in Xinjiang, which was consistent with the gene flow between West and East Eurasian. J2a has a high frequency in Qinghai, situated within the Hexi corridor, which might imply the impact of the Silk Road connecting China with the West (Yang et al. 2008; Shou et al. 2010). These findings elucidated the intricate interaction between the Han Chinese and ethnolinguistically diverse East Asian groups, shedding light on gene flow dynamics between West and East Eurasia.

**Figure 6.**
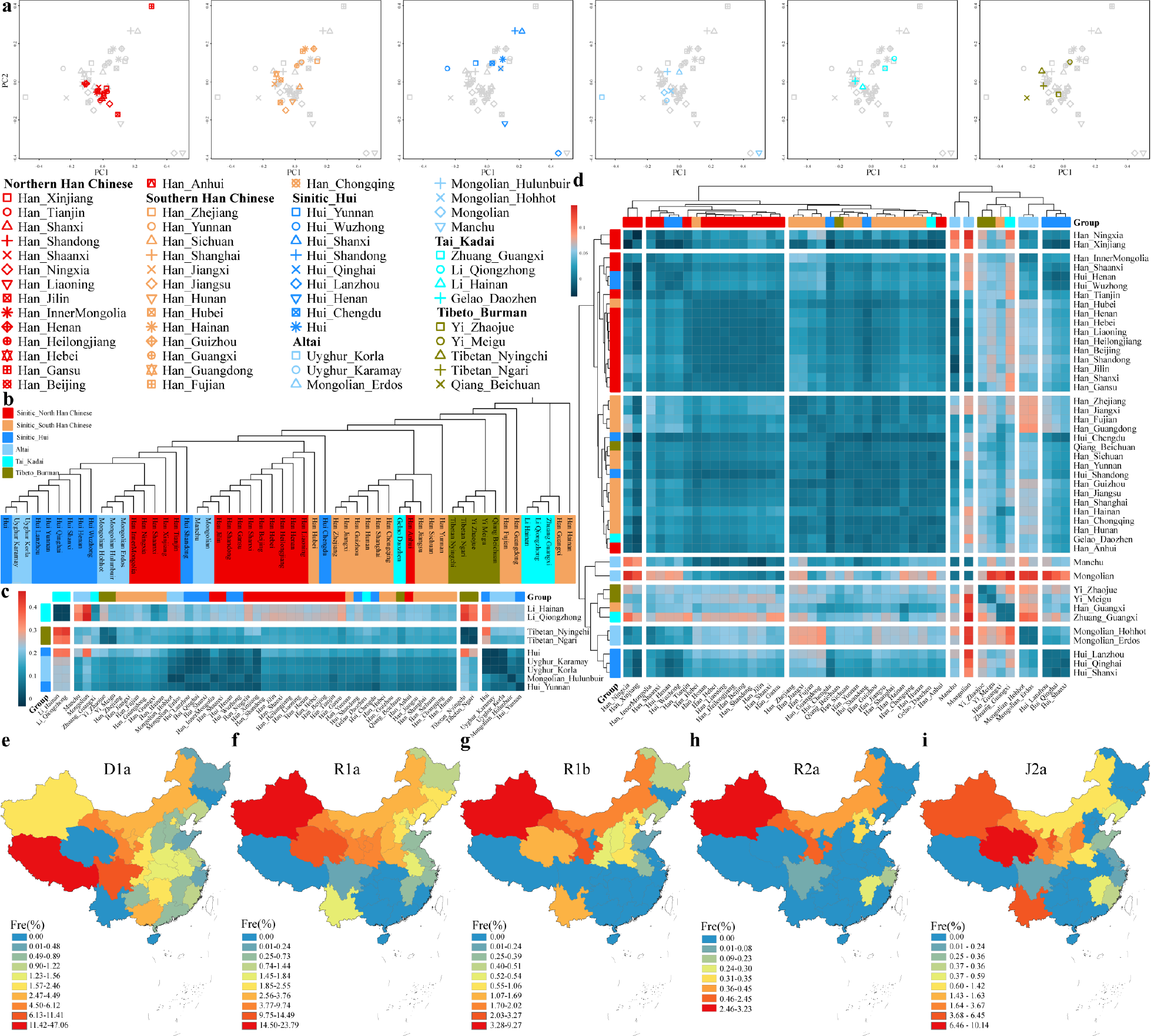
The genetic relationship among East Asians inferred from Y chromosome variations. **(a)** PCA inferred from the haplogroup frequency matrix among 54 ethnolinguistically diverse populations in East Asia. **(b)** The neighbor-joining tree among 54 ethnolinguistically diverse populations in East Asia. **(c-d)** The heatmap of pairwise F_ST_ genetic distances among 54 ethnolinguistically diverse populations in East Asia with different legend values. **(e-i)** The haplogroup frequency among 54 ethnolinguistically diverse populations in East Asia.

## Discussion

The proliferation of large-scale genetic studies focusing on autosomal DNA advanced our understanding of fine-scale genetic structure and demographic history (Bai et al. 2018; Liu et al. 2021; Ma et al. 2021; Chen et al. 2022; Pan et al. 2022). Moreover, Y-chromosome and mtDNA variations provided a unique insight into sex-specific genetic history (Wen et al. 2004a; Kutanan et al. 2019; Li et al. 2019a; Pinotti et al. 2019; Kutanan et al. 2020; Macholdt et al. 2020; Karmin et al. 2022). However, these researches focusing on the Y-chromosome in East Asia tended to concentrate on the phylogenetic variations, which more or less overlooked the investigation of paternal genetic structure and corresponding factors in the formation of the Han Chinese. Here, to strengthen our understanding of the fine-scale demographical history of the Han Chinese, we launched the YanHuang cohort and presented the repertoires of Y-chromosome variations from 5,020 Han Chinese belonging to 29 different administrative provinces sequencing via our developed YHseqY3000 panel. The high value of haplotype and haplogroup-based parameters indicated the strong performance of our developed panel in haplogroup allocation and variation capture. Moreover, we presented one of the most extensive HFS of the Han Chinese, in which O2a, O1b, O1a and C2b accounted for most of the haplogroup composition. In particular, we found a significant paternal genetic differentiation corresponding to the geographic boundaries of the Qinling-Huaihe and Nanling Mountains and population admixture among the western and eastern Eurasian populations, as well as the northern and southern East Asian populations. Here, we will discuss the isolation-enhanced and admixture-introduced genetic differentiation contributed to the formation mechanism of the Han Chinese and summarized this observed pattern as the WDMSA model.

### Isolation-enhanced genetic differentiation on the paternal genetic structure

Our study observed the North-South genetic divergence in the Han Chinese, consistent with previous studies based on autosomal DNA and mtDNA (Wen et al. 2004a; Chen et al. 2009). At the same time, our results further separated the Han Chinese into three subgroups, including North/South/Lingnan Han Chinese, corresponding to the geographical boundaries **(Figure 2a)**. Abundant previous evidence based on the autosomal variations revealed four subgroups of the Han Chinese, including North/South/Central/Lingnan Han, in which the Central Han showed a close pairwise genetic distance with North Han than with South Han and Lingnan Han (Cong et al. 2022). However, our study showed that the Central Han showed a close genetic distance with South Han (0.0022, *P* < 0.01) instead of North Han (0.0089, *P* < 0.01) **(Table S5)**. The catalog of genetic structure implied different genetic histories revealed by the Y-chromosome and autosomal variations.

AMOVA-based results inferred that the largest among-group variation (1.14%, *P* < 0.001) was observed in the grouping according to geographical boundaries **(Table 1)**, which further confirmed the pattern of genetic stratification. In addition, northern Han Chinese performed more homogeneous, whereas southern Han Chinese were more diverse than their northern counterpart **(Table 1)**, which might result from the prehistoric migration and interaction between southern Han Chinese and southern Indigenous groups (Su et al. 1999; Wen et al. 2004a). The Qinling-Huaihe Line is related to the division line of hydrology and climatology and roughly corresponds to the line between Millet-based farming in the YRB and Rice-based farming in the Yangtze River Basin (Li et al. 2012). The Nanling Mountain serves as a geographical barrier between the Yangtze River Basin and the Zhujiang River Basin. This might lead to the among-group variation observed among the Yangtze River, Yellow River and Zhujiang River populations, thus exhibiting a relatively elevated degree of differentiation.

Otherwise, the correlation between language dialects and autosomal DNA had been presented in previous studies (Qiu et al. 2022) but rare for the mtDNA (Li et al. 2019b). In our study, we noticed that the Han Chinese speaking Mandarin presented genetic differences with other dialect groups **(Figure 2c)**, possibly due to the patrilocality residence customs in East Asia (Wang and Li 2013). Importantly, we only observed the genetic divergence between Mandarin speakers and other dialect speakers. Considering the observed paternal genetic structure and genetic divergence, the isolation-enhanced genetic differentiation presented by geography played an essential role in shaping the paternal structure of the Han Chinese. In addition, the river basin is always related to agricultural traditions, which are strongly related to population expansion and have been proven to influence the maternal genetic landscape of the Han Chinese. The future whole-genome sequencing on the Y-chromosome will provide more clues for whether agricultural traditions have affected the paternal landscape of the Han Chinese.

### Admixture-introduced genetic differentiation on the paternal genetic structure

Population admixture is one of the most essential driving forces influencing genomic diversity, which can decrease the between-population differentiation and increase the within-population genetic diversity. In our study, the evidence regarding Fisher’s exact test and mantel test that elaborated the differentiation of paternal genetic structure was accounted for O1a, O1b, C2b, C2a, N1a, R1a, R1b and J2a **(Figure 2d and Table S7)**, which also correlated with the four ancestral components simulated by ADMIXTURE **(Figure 4c)**. Therefore, the observed correlation suggested that these haplogroups contributing to the genetic differentiation of the Han Chinese were introduced by admixture. Of note, due to the sample bias and paucity of the ancient DNA in East Asia, the correlation analysis might increase the false positive rate, such as the D1a and AWES-related populations **(Figure 4c)**. However, most of the significant correlations can be verified from the archaeogenetics. We can observe these haplogroups arising in corresponding ancient populations, but it is difficult to determine the concrete time of admixture. In particular, the O2a haplogroup frequently emerged in ANEA-related populations and took account of the majority of the paternal genetic landscape and did not contribute to the genetic divergence, which might indicate that the demic diffusion of the Han Chinese from north to south was mainly driven by ancient populations carrying O2a haplogroup.

The formation of the Han Chinese can be viewed as a tapestry interweaved by admixture events. According to the historical record, the Han Chinese were generally thought to be descended from the ancient Huaxia tribe located in the upper and middle YRB. About six kya, the proto-Han Chinese were divided from the proto-ST people and migrated to the south and east of East Asia. Long-term migration, interaction and admixture with surrounding populations led to the largest population size and wide distribution of the Han Chinese in China. Consequently, the WDMSA model within the Han Chinese is much simplified. However, we identified multiple ancestral sources from different Eurasian regions that contributed the genetic legacy to the formation of Han Chinese’s complex paternal gene pool, which was consistent with our proposed admixture-by-admixture models with some weakly differentiated and connected ancestral sources. The analysis of autosomal variations of ancient populations also confirmed this finding. For example, Fu et al. revealed the gene flow between northern and southern China since the Neolithic, as the genetic affinity was higher in the present than in the past (Yang et al. 2020). Wang et al. also found that ancient farmers from Taiwan received the gene pool from northern ancestry in the Neolithic (Wang et al. 2021a). Jeong et al. identified the gene flow from the western pastoralists to the eastern Eurasian cal. 3000 BCE (Jeong et al. 2020). The WDMSA model proposed here provided a general insight into the Han Chinese’s formation, but the Han Chinese’s dynamic migration and admixture history are multi-wave and multi-level. In the future, the broader coverage of sampling and higher resolution sequence, together with more paleogenomic studies, are expected to provide a more detailed evolutionary history of the Han Chinese.

### Limitations

Y-chromosome complex tandem regions, especially in the telomere, centromere and segmental duplication regions, were the hardy-measured sequenced regions in the past years (Zhou et al. 2023). Our presentation focused on the high-confidence lineage markers in the male-specific regions of the Y-chromosome based on the capturing target sequencing to present the full landscape of paternal lineages of Han Chinese populations. Recent advantages in second and third-generation long-read sequencing (Oxford Nanopore Technologies and PacBio HiFi) and statistical assembly and genotyping innovations provided further insights for the landscape of the complete sequence and revolutionary features of pseudoautosomal regions (PARs), X-degenerate regions (XDR), X-degenerate regions (XDR), ampliconic palindromic regions, q-arm heterochromatin and centromeric satellites of a complete Y chromosome (Hallast et al. 2023; Rhie et al. 2023). Thus, high-coverage complete whole-genome Y-chromosome sequences from genetically different populations in our YHC, 10K_CPGDP and other Chinese cohorts were needed to provide deep insights into the formation of Han Chinese. Besides, Samping bias exists here, especially in the high-altitude Tibetan Plateau. Further anthropologically informed sampling across a wider region of Han Chinese and minority ethnic groups will also be needed to eliminate the interactions among ethnolinguistically underrepresented populations.

## Conclusion

We reported the largest paternal genomic resources from the YanHuang cohort and presented one large-scale and extensive paternal genomic data of the Han Chinese by genotyping 5,020 Han Chinese individuals from 29 different geographical populations. We identified fine-scale paternal genetic structures, including North, South and Lingnan Han along the Qinling-Haihe and Nanling mountains and multiple and complex founding lineages in geographically different Han Chinese, whose frequency distribution possessed intense latitude-dependent gradient changes. Most importantly, we identified two major evolutionary forces, including isolation-enhanced and admixture-introduced genetic differentiation, which obviously shaped the paternal genetic structure. Meanwhile, we finally proposed the new admixture-by-admixture WDMSA model to deliver the multiple, complex and multi-layer ancestral sources related to local ancient millet and rice farmers and neighboring Siberian hunter-gatherers and herders contributed to the genomic information of geographically different Han Chinese. In summary, we provided new insights into the paternal genetic structure of the Han Chinese and the influencing factors of their formation based on the newly collected paternal genomic sources and reported a most plausible complex admixture model to illuminate the evolutionary history of the Han Chinese.

## MATERIALS AND METHODS

### Studied populations

We collected peripheral blood from 5,020 unrelated males distributed in 29 different administrative provinces (**Figure 1a**), including Anhui (N = 236), Beijing (N = 133), Chongqing (N = 99), Fujian (N = 142), Gansu (N = 65), Guangdong (N = 387), Guangxi (N = 89), Guizhou (N = 64), Hainan (N = 11), Hebei (N = 230), Heilongjiang (N = 92), Henan (N = 253), Hubei (N = 183), Hunan (N = 198), InnerMongolia (N = 41), Jiangsu (N = 499), Jiangxi (N = 196), Jilin (N = 69), Liaoning (N = 161), Ningxia (N = 12), Shaanxi (N = 158), Shandong (N = 594), Shanghai (N = 137), Shanxi (N = 158), Sichuan (N = 342), Tianjin (N = 69), Xinjiang (N = 10), Yunnan (N = 62) and Zhejiang (N = 331). Each donor provided the informed consent. Ethical approval was provided by the Medical Ethics Committee of West China Hospital of Sichuan University (2023-1321). Our experiment obeyed the recommendations and regulations of our institute and national guidelines of the standard of the Declaration of Helsinki (Association 2013). Genomic DNA was extracted using the QIAamp DNA Mini Kit (QIAGEN, Germany). We used the Qubit dsDNA HS Assay Kit to quantify DNA concentrations based on the standard protocol on a Qubit 3.0 fluorometer (Thermo Fisher Scientific). Then, DNA samples were stored at – 20°C until amplification.

### Genotyping and quality control

We sequenced the targeted Y-chromosome regions in 5,020 samples using the YHseqY3000 panel on the Illumina platform (Illumina, San Diego, CA, USA) and used the BWA v.0.7.13 (Li and Durbin 2009) to map the raw sequencing reads to the human reference genome GRCh37. The Quality control was conducted by the PLINK v1.90b6.26 64-bit (2 Apr 2022) with two parameters (-geno: 0.1 and -mind: 0.1) (Chang et al. 2015).

### Haplogroup allocation

First, we classify the haplogroup with our developed forensic phylogenetic tree using our in-house pipeline. Consistent with previous research on the haplogroup Nomenclature, the haplogroup for each sample was allocated by the python package of hGrpr2 in HaploGrouper based on the ISOGG Y-DNA Haplogroup Tree 2019–2020 version 15.73 (Jagadeesan et al. 2021). In addition, we used Y-LineageTracker to classify the haplogroup for comparing the consistency of haplogroup allocation (Chen et al. 2021). Detailed results are listed in **Table S2**.

### The performance of our developed panel

We used Arlequin 3.5.1.3 to calculate the haplotype frequency of 29 populations, respectively. We calculated the haplotype/haplogroup-based forensic parameter with the following formulas. Haplotype diversity (HD) was calculated based on the following formula from Nei and Tajima: HD = *n* (1 − Σ*pi*^2^)/(n-1), where *n* is the mean of the total number of observed haplotypes and *pi* is the mean of the frequency of the i-th haplotype. Discrimination capacity (DC) was calculated as the ratio between the number of observed haplotypes and the number of total haplotypes. Match probability (MP) was computed as the sum of squared haplotype frequencies: MP = Σ*pi*^2^.

## Statistical Analyses

### Y-chromosome-based analyses

#### Genetic diversity

We used the Python package ClusterHaplogroup in Y-LineageTracker to calculate the haplogroup frequency of 29 populations from different provinces. To prevent the potential bias from the small sample size (< 30), we removed the samples from Xinjiang, Ningxia and Hainan for further analysis. Using the R package, we generated the HFS within the third level. We used ArcGIS software to present the geographic distribution of the haplogroup frequency and conducted the spatial autocorrelation analysis with the above results. Moran’s I index can dissect whether the haplogroup distribution has clustering properties in spatial geographical distribution. Getis-Ord Gi* can distinguish the HotSpot and ColdSpot related to a cluster region with high and low values, respectively. The HotSpot generally corresponds to the center of diffusion of the haplogroup.

#### Genetic relationship and population structure

We used the Python package of ClusterHaplogroup in Y-LineageTracker to perform PCA based on the haplogroup frequency at the fourth level. We also used Arlequin 3.5.1.3 to calculate the pairwise genetic distances (F_ST_) within 26 populations and then carried out a nonparametric MDS analysis using the R package (Excoffier and Lischer 2010). We conducted the correlation analysis and mantel test using the R package. We constructed an unrooted neighbor-joining tree based on genetic distances (F_ST_) among 26 populations using MEGA 7 (Kumar et al. 2016). We also generated the genetic distances (F_ST_) matrix using the R package among 26 populations. We performed the AMOVA analysis using the Arlequin 3.5.1.3. We performed the mantel test to explore the model of isolate by distance, which meant the relationship between the genetic distance and the geographic distance using the R package with 10000 rounds of permutation.

#### Phylogeny analysis and network analysis

We generated the Newick file of the phylogenetic tree of all samples using the Python package of PhyloHaplogroup in Y-LineageTracker with the maximum parsimony method. Then, we import the Newick file into the website of iTOL: Interactive Tree of Life to annotate and embellish. We used the Python package of fasta_to_nexus_Main.py (fasta_nexus_converter/Main.py at master· rubenAlbuquerque/fasta_nexus_converter· GitHub) to convert Fasta file to Nexus files. Then, we used popART to generate the Median-Joining Network (Bandelt et al. 1999; Leigh et al. 2015).

### Autosome-based analyses

#### Model-based ADMIXTURE analysis

We used PLINK v.1.90 to prune the original dataset with strong linkage disequilibrium SNPs based on the parameters of --indep-pairwise 200 25 0.4 (Chang et al. 2015). Then, we applied the unsupervised ADMIXTURE v.1.3.0 analysis based on the clustering algorithm of maximum likelihood (Alexander et al. 2009) to explore the genetic structure and identify ancestral sources in the context of different regions in China at the autosomal chromosome level. The model-based ADMIXTURE analysis included 1,132 individuals from 32 Han Chinese populations from different administrative provinces and 89 ancient and modern reference populations included in our merged 10K_CPGDP database. On this basis, we ran ADMIXTURE for 100 iterations with the number of ancestral sources from K = 2 to K = 20 through the default parameters. Besides, we performed 10-fold cross-validation (--cv = 10) using different random seeds to determine the optimal K value (Feng et al. 2018), which depended on the lowest cross-validation error and the highest log-likelihood. We used ArcMap 10.8 to visualize the frequency distribution of four ancestral components among 32 populations of the Han Chinese from different administrative provinces. We then correlated the four ancestral components with the paternal lineage frequency and visualized the result of correlation analysis using the R package.

#### ADMIXTOOLS2 analysis

For the ADMIXTOOLS2 analysis, we integrated data from 32 Han Chinese populations along with continentally representative modern ancient source populations (Maier et al. 2023), including Yoruba, Russia_Afanasievo, China_SEastAsia_Island_EN, Mongolia_North_N, China_YR_LN and GaoHuaHua. The function *find_graphs* in the R package ADMIXTOOLS2 was setting 0 to 2 admixture events with 50 replicates for each Han Chinese population. This approach leveraged *f*-statistics and the optimal model fit is shown in **Figure S7**.

## Declarations

### Ethics approval and consent to participate

The Medical Ethics Committee of West China Hospital of Sichuan University approved this study. This study was conducted in accordance with the principles of the Helsinki Declaration.

### Consent for publication

Not applicable.

### Data Availability

All haplogroup information was submitted in the supplementary materials. We followed the regulations of the Ministry of Science and Technology of the People’s Republic of China. The raw genotype data required controlled access. Further requests for access to data can be directed to Guanglin He and Mengge Wang.

### Author Contributions

Z.W., G.H., M.W., S.N. and C.L. conceived and supervised the project. G.H. and M.W. collected the samples. Z.W., Y.H., G.H. and M.W. extracted the genomic DNA and performed the genome sequencing. Z.W., G.H., M.W. and S.D. did variant calling. Z.W., B.J., L.L., Y.L., X.J., M.W., Y.H., S.D., R.T., K.L., L.H., J.C., X.L.,Q.Y., Y.S., Q.S., Y.H., H.S., J.Z., H.Y., L.Y., J.Y., Y.C., K.Z., H.D., J.Y., B.Z., S.N., C.L., G.H. performed population genetic analysis. Z.W., G.H. and M.W. drafted the manuscript. Z.W., G.H., M.W., S.N. and C.L. revised the manuscript.

## Acknowledgments

We thank all the volunteers who participated in this project.

## ^†^Full author lists of 10K_CPGDP consortium

Mengge Wang^1,2,4,5^, Haibing Yuan^2^, Lanhai Wei^9^, Renkuan Tang^11^, Liping Hu^3^, Yuguo Huang^1^, Hongbing Yao^10^, Libing Yun^13^, Junbao Yang^14^, Yan Cai^15^, Hong Deng^3^, Jiangwei Yan^12^, Bofeng Zhu^16,17^, Shengjie Nie^2^, Chao Liu^5,15,18^, Guanglin He^1,2^

## REFERENCES

Alexander DH, Novembre J, Lange K. 2009. Fast model-based estimation of ancestry in unrelated individuals. Genome Res 19: 1655–1664.

Association WM. 2013. World Medical Association Declaration of Helsinki: ethical principles for medical research involving human subjects. Jama 310: 2191–2194.

Bai H, Guo X, Narisu N, Lan T, Wu Q, Xing Y, Zhang Y, Bond SR, Pei Z, Zhang Y et al. 2018. Whole-genome sequencing of 175 Mongolians uncovers population-specific genetic architecture and gene flow throughout North and East Asia. Nat Genet 50: 1696–1704.

Bandelt HJ, Forster P, Rohl A. 1999. Median-joining networks for inferring intraspecific phylogenies. Mol Biol Evol 16: 37–48.

Bellwood P. 2005. Examining the farming/language dispersal hypothesis in the East Asian context. In The Peopling of East Asia, pp. 41–54. Routledge.

Cao Y, Li L, Xu M, Feng Z, Sun X, Lu J, Xu Y, Du P, Wang T, Hu R et al. 2020. The ChinaMAP analytics of deep whole genome sequences in 10,588 individuals. Cell Res 30: 717–731.

Cavalli-Sforza LL. 1998. The Chinese human genome diversity project. Proc Natl Acad Sci U S A 95: 11501–11503.

Cavalli-Sforza LL, Menozzi P, Piazza A. 1994. The history and geography of human genes. Princeton university press.

Chang CC, Chow CC, Tellier LC, Vattikuti S, Purcell SM, Lee JJ. 2015. Second-generation PLINK: rising to the challenge of larger and richer datasets. Gigascience 4: 7.

Chen H, Lin R, Lu Y, Zhang R, Gao Y, He Y, Xu S. 2022. Tracing Bai-Yue Ancestry in Aboriginal Li People on Hainan Island. Mol Biol Evol 39: msac210.

Chen H, Lu Y, Lu D, Xu S. 2021. Y-LineageTracker: a high-throughput analysis framework for Y-chromosomal next-generation sequencing data. BMC Bioinformatics 22: 114.

Chen J, Zheng H, Bei JX, Sun L, Jia WH, Li T, Zhang F, Seielstad M, Zeng YX, Zhang X et al. 2009. Genetic structure of the Han Chinese population revealed by genome-wide SNP variation. Am J Hum Genet 85: 775–785.

Chen P, Wu J, Luo L, Gao H, Wang M, Zou X, Li Y, Chen G, Luo H, Yu L et al. 2019. Population Genetic Analysis of Modern and Ancient DNA Variations Yields New Insights Into the Formation, Genetic Structure, and Phylogenetic Relationship of Northern Han Chinese. Front Genet 10: 1045.

Cheng S, Xu Z, Bian S, Chen X, Shi Y, Li Y, Duan Y, Liu Y, Lin J, Jiang Y. 2023. The STROMICS genome study: deep whole-genome sequencing and analysis of 10K Chinese patients with ischemic stroke reveal complex genetic and phenotypic interplay. Cell Discovery 9: 75.

Chiang CW, Mangul S, Robles C, Sankararaman S. 2018. A comprehensive map of genetic variation in the world’s largest ethnic group—Han Chinese. Mol Biol Evol 35: 2736–2750.

Cong PK, Bai WY, Li JC, Yang MY, Khederzadeh S, Gai SR, Li N, Liu YH, Yu SH, Zhao WW et al. 2022. Genomic analyses of 10,376 individuals in the Westlake BioBank for Chinese (WBBC) pilot project. Nat Commun 13: 2939.

Consortium HP-AS, Abdulla MA, Ahmed I, Assawamakin A, Bhak J, Brahmachari SK, Calacal GC, Chaurasia A, Chen CH, Chen J et al. 2009. Mapping human genetic diversity in Asia. Science 326: 1541–1545.

Damgaard PB, Marchi N, Rasmussen S, Peyrot M, Renaud G, Korneliussen T, Moreno-Mayar JV, Pedersen MW, Goldberg A, Usmanova E et al. 2018. 137 ancient human genomes from across the Eurasian steppes. Nature 557: 369–374.

Diamond J, Bellwood P. 2003. Farmers and their languages: the first expansions. Science 300: 597–603.

Excoffier L, Lischer HE. 2010. Arlequin suite ver 3.5: a new series of programs to perform population genetics analyses under Linux and Windows. Mol Ecol Resour 10: 564–567.

Feng Q, Lu D, Xu S. 2018. AncestryPainter: a graphic program for displaying ancestry composition of populations and individuals. Genomics, Proteomics & Bioinformatics 16: 382–385.

Ge J, Wu S, Chao S. 1997. Zhongguo yimin shi (The migration history of China). Fujian People’s Publishing House, Fuzhou, China.

GenomeAsia KC. 2019. The GenomeAsia 100K Project enables genetic discoveries across Asia. Nature 576: 106–111.

Hallast P, Ebert P, Loftus M, Yilmaz F, Audano PA, Logsdon GA, Bonder MJ, Zhou W, Hops W, Kim K et al. 2023. Assembly of 43 human Y chromosomes reveals extensive complexity and variation. Nature 621: 355–364.

He G, Wang M, Miao L, Chen J, Zhao J, Sun Q, Duan S, Wang Z, Xu X, Sun Y et al. 2023a. Multiple founding paternal lineages inferred from the newly-developed 639-plex Y-SNP panel suggested the complex admixture and migration history of Chinese people. Human genomics 17.

He G, Wang Z, Guo J, Wang M, Zou X, Tang R, Liu J, Zhang H, Li Y, Hu R et al. 2020. Inferring the population history of Tai-Kadai-speaking people and southernmost Han Chinese on Hainan Island by genome-wide array genotyping. Eur J Hum Genet 28: 1111–1123.

He G, Yao H, Sun Q, Duan S, Tang R, Chen J, Wang Z, Sun Y, Li X, Wang S et al. 2023b. Whole-genome sequencing of ethnolinguistic diverse northwestern Chinese Hexi Corridor people from the 10K_CPGDP project suggested the differentiated East-West genetic admixture along the Silk Road and their biological adaptations. bioRxiv doi:10.1101/2023.02.26.530053: 2023.2002.2026.530053.

He GL, Li YX, Zou X, Yeh HY, Tang RK, Wang PX, Bai JY, Yang XM, Wang Z, Guo JX et al. 2022. Northern gene flow into southeastern East Asians inferred from genome-wide array genotyping. Journal of Systematics and Evolution 61: 179–197.

He GL, Wang MG, Li YX, Zou X, Yeh HY, Tang RK, Yang XM, Wang Z, Guo JX, Luo T et al. 2021. Fine-scale north-to-south genetic admixture profile in Shaanxi Han Chinese revealed by genome-wide demographic history reconstruction. Journal of Systematics and Evolution 60: 955–972.

Jagadeesan A, Ebenesersdottir SS, Guethmundsdottir VB, Thordardottir EL, Moore KHS, Helgason A. 2021. HaploGrouper: a generalized approach to haplogroup classification. Bioinformatics 37: 570–572.

Jeong C, Ozga AT, Witonsky DB, Malmstrom H, Edlund H, Hofman CA, Hagan RW, Jakobsson M, Lewis CM, Aldenderfer MS et al. 2016. Long-term genetic stability and a high-altitude East Asian origin for the peoples of the high valleys of the Himalayan arc. Proc Natl Acad Sci U S A 113: 7485–7490.

Jeong C, Wang K, Wilkin S, Taylor WTT, Miller BK, Bemmann JH, Stahl R, Chiovelli C, Knolle F, Ulziibayar S et al. 2020. A Dynamic 6,000-Year Genetic History of Eurasia’s Eastern Steppe. Cell 183: 890–904 e829.

Jin L, Su B. 2000. Natives or immigrants: modern human origin in east Asia. Nat Rev Genet 1: 126–133.

Karafet TM, Mendez FL, Meilerman MB, Underhill PA, Zegura SL, Hammer MF. 2008. New binary polymorphisms reshape and increase resolution of the human Y chromosomal haplogroup tree. Genome Res 18: 830–838.

Karmin M, Flores R, Saag L, Hudjashov G, Brucato N, Crenna-Darusallam C, Larena M, Endicott PL, Jakobsson M, Lansing JS et al. 2022. Episodes of Diversification and Isolation in Island Southeast Asian and Near Oceanian Male Lineages. Mol Biol Evol 39.

Kumar S, Stecher G, Tamura K. 2016. MEGA7: Molecular Evolutionary Genetics Analysis Version 7.0 for Bigger Datasets. Mol Biol Evol 33: 1870–1874.

Kumar V, Wang W, Zhang J, Wang Y, Ruan Q, Yu J, Wu X, Hu X, Wu X, Guo W et al. 2022. Bronze and Iron Age population movements underlie Xinjiang population history. Science 376: 62–69.

Kutanan W, Kampuansai J, Srikummool M, Brunelli A, Ghirotto S, Arias L, Macholdt E, Hubner A, Schroder R, Stoneking M. 2019. Contrasting Paternal and Maternal Genetic Histories of Thai and Lao Populations. Mol Biol Evol 36: 1490–1506.

Kutanan W, Shoocongdej R, Srikummool M, Hubner A, Suttipai T, Srithawong S, Kampuansai J, Stoneking M. 2020. Cultural variation impacts paternal and maternal genetic lineages of the Hmong-Mien and Sino-Tibetan groups from Thailand. Eur J Hum Genet 28: 1563–1579.

Lang M, Liu H, Song F, Qiao X, Ye Y, Ren H, Li J, Huang J, Xie M, Chen S et al. 2019. Forensic characteristics and genetic analysis of both 27 Y-STRs and 143 Y-SNPs in Eastern Han Chinese population. Forensic Sci Int Genet 42: e13–e20.

Lazaridis I Patterson N Mittnik A Renaud G Mallick S Kirsanow K Sudmant PH Schraiber JG Castellano S Lipson M et al. 2014. Ancient human genomes suggest three ancestral populations for present-day Europeans. Nature 513: 409–413.

Leigh JW, Bryant D, Nakagawa S. 2015. popart: full-feature software for haplotype network construction. Methods in Ecology and Evolution 6: 1110–1116.

Li H, Durbin R. 2009. Fast and accurate short read alignment with Burrows-Wheeler transform. Bioinformatics 25: 1754–1760.

Li H, Huang Y, Mustavich LF, Zhang F, Tan JZ, Wang LE, Qian J, Gao MH, Jin L. 2007. Y chromosomes of prehistoric people along the Yangtze River. Hum Genet 122: 383–388.

Li S, Yan J, Wan J. 2012. The characteristics of temperature change in Qinling Mountains. Sci Geogr Sin 32: 853–858.

Li Y-C, Gao Z-L, Liu K-J, Tian J-Y, Yang B-Y, Rahman ZU, Yang L-Q, Zhang S-H, Li C-T, Achilli A. 2023. Mitogenome evidence shows two radiation events and dispersals of matrilineal ancestry from northern coastal China to the Americas and Japan. Cell Reports.

Li YC, Tian JY, Liu FW, Yang BY, Gu KS, Rahman ZU, Yang LQ, Chen FH, Dong GH, Kong QP. 2019a. Neolithic millet farmers contributed to the permanent settlement of the Tibetan Plateau by adopting barley agriculture. Natl Sci Rev 6: 1005–1013.

Li YC, Ye WJ, Jiang CG, Zeng Z, Tian JY, Yang LQ, Liu KJ, Kong QP. 2019b. River Valleys Shaped the Maternal Genetic Landscape of Han Chinese. Mol Biol Evol 36: 1643–1652.

Liang Z, White MJ. 1996. Internal migration in China, 1950–1988. Demography 33: 375–384.

Lipson M, Cheronet O, Mallick S, Rohland N, Oxenham M, Pietrusewsky M, Pryce TO, Willis A, Matsumura H, Buckley H et al. 2018. Ancient genomes document multiple waves of migration in Southeast Asian prehistory. Science 361: 92–95.

Liu CC, Witonsky D, Gosling A, Lee JH, Ringbauer H, Hagan R, Patel N, Stahl R, Novembre J, Aldenderfer M et al. 2022a. Ancient genomes from the Himalayas illuminate the genetic history of Tibetans and their Tibeto-Burman speaking neighbors. Nat Commun 13: 1203.

Liu L, Chen J, Wang J, Zhao Y, Chen X. 2022b. Archaeological evidence for initial migration of Neolithic Proto Sino-Tibetan speakers from Yellow River valley to Tibetan Plateau. Proc Natl Acad Sci U S A 119: e2212006119.

Liu S, Huang S, Chen F, Zhao L, Yuan Y, Francis SS, Fang L, Li Z, Lin L, Liu R. 2018. Genomic analyses from non-invasive prenatal testing reveal genetic associations, patterns of viral infections, and Chinese population history. Cell 175: 347–359. e314.

Liu Y, Xie J, Wang M, Liu C, Zhu J, Zou X, Li W, Wang L, Leng C, Xu Q et al. 2021. Genomic Insights Into the Population History and Biological Adaptation of Southwestern Chinese Hmong-Mien People. Front Genet 12: 815160.

Ma X, Yang W, Gao Y, Pan Y, Lu Y, Chen H, Lu D, Xu S. 2021. Genetic Origins and Sex-Biased Admixture of the Huis. Mol Biol Evol 38: 3804–3819.

Macholdt E, Arias L, Duong NT, Ton ND, Van Phong N, Schroder R, Pakendorf B, Van Hai N, Stoneking M. 2020. The paternal and maternal genetic history of Vietnamese populations. Eur J Hum Genet 28: 636–645.

Maier R, Flegontov P, Flegontova O, Isildak U, Changmai P, Reich D. 2023. On the limits of fitting complex models of population history to f-statistics. eLife 12: e85492.

Mao X, Zhang H, Qiao S, Liu Y, Chang F, Xie P, Zhang M, Wang T, Li M, Cao P et al. 2021. The deep population history of northern East Asia from the Late Pleistocene to the Holocene. Cell 184: 3256–3266 e3213.

McColl H, Racimo F, Vinner L, Demeter F, Gakuhari T, Moreno-Mayar JV, van Driem G, Gram Wilken U, Seguin-Orlando A, de la Fuente Castro C et al. 2018. The prehistoric peopling of Southeast Asia. Science 361: 88–92.

Narasimhan VM Patterson N Moorjani P Rohland N Bernardos R Mallick S Lazaridis I Nakatsuka N Olalde I Lipson M et al. 2019. The formation of human populations in South and Central Asia. Science 365: eaat7487.

Ning C, Li T, Wang K, Zhang F, Li T, Wu X, Gao S, Zhang Q, Zhang H, Hudson MJ et al. 2020. Ancient genomes from northern China suggest links between subsistence changes and human migration. Nat Commun 11: 2700.

Pan Y, Zhang C, Lu Y, Ning Z, Lu D, Gao Y, Zhao X, Yang Y, Guan Y, Mamatyusupu D et al. 2022. Genomic diversity and post-admixture adaptation in the Uyghurs. Natl Sci Rev 9: nwab124.

Pinotti T, Bergstrom A, Geppert M, Bawn M, Ohasi D, Shi W, Lacerda DR, Solli A, Norstedt J, Reed K et al. 2019. Y Chromosome Sequences Reveal a Short Beringian Standstill, Rapid Expansion, and early Population structure of Native American Founders. Curr Biol 29: 149–157 e143.

Qi X, Cui C, Peng Y, Zhang X, Yang Z, Zhong H, Zhang H, Xiang K, Cao X, Wang Y et al. 2013. Genetic evidence of paleolithic colonization and neolithic expansion of modern humans on the tibetan plateau. Mol Biol Evol 30: 1761–1778.

Qiu X, Huang S, Huang M, Liu S, Wang C, He J, Kuang Y, Lu J, Gu Y, Xia X et al. 2022. Whole genome sequencing and analysis of 4,053 individuals in trios and mother-infant duos from the Born in Guangzhou Cohort Study. doi:10.21203/rs.3.rs-1732885/v1.

Ragsdale AP, Weaver TD, Atkinson EG, Hoal EG, Moller M, Henn BM, Gravel S. 2023. A weakly structured stem for human origins in Africa. Nature 617: 755–763.

Reich D, Thangaraj K, Patterson N, Price AL, Singh L. 2009. Reconstructing Indian population history. Nature 461: 489–494.

Rhie A, Nurk S, Cechova M, Hoyt SJ, Taylor DJ, Altemose N, Hook PW, Koren S, Rautiainen M, Alexandrov IA et al. 2023. The complete sequence of a human Y chromosome. Nature 621: 344–354.

Shi H, Zhong H, Peng Y, Dong YL, Qi XB, Zhang F, Liu LF, Tan SJ, Ma RZ, Xiao CJ et al. 2008. Y chromosome evidence of earliest modern human settlement in East Asia and multiple origins of Tibetan and Japanese populations. BMC Biol 6: 45.

Shou WH, Qiao EF, Wei CY, Dong YL, Tan SJ, Shi H, Tang WR, Xiao CJ. 2010. Y-chromosome distributions among populations in Northwest China identify significant contribution from Central Asian pastoralists and lesser influence of western Eurasians. J Hum Genet 55: 314–322.

Song F, Song M, Luo H, Xie M, Wang X, Dai H, Hou Y. 2021. Paternal genetic structure of Kyrgyz ethnic group in China revealed by high-resolution Y-chromosome STRs and SNPs. Electrophoresis 42: 1892–1899.

Song M, Wang Z, Lyu Q, Ying J, Wu Q, Jiang L, Wang F, Zhou Y, Song F, Luo H et al. 2022. Paternal genetic structure of the Qiang ethnic group in China revealed by high-resolution Y-chromosome STRs and SNPs. Forensic Sci Int Genet 61: 102774.

Song M, Wang Z, Zhang Y, Zhao C, Lang M, Xie M, Qian X, Wang M, Hou Y. 2019. Forensic characteristics and phylogenetic analysis of both Y-STR and Y-SNP in the Li and Han ethnic groups from Hainan Island of China. Forensic Sci Int Genet 39: e14–e20.

Su B, Xiao J, Underhill P, Deka R, Zhang W, Akey J, Huang W, Shen D, Lu D, Luo J et al. 1999. Y-Chromosome evidence for a northward migration of modern humans into Eastern Asia during the last Ice Age. Am J Hum Genet 65: 1718–1724.

Sun J, Li YX, Ma PC, Yan S, Cheng HZ, Fan ZQ, Deng XH, Ru K, Wang CC, Chen G et al. 2021. Shared paternal ancestry of Han, Tai-Kadai-speaking, and Austronesian-speaking populations as revealed by the high resolution phylogeny of O1a-M119 and distribution of its sub-lineages within China. Am J Phys Anthropol 174: 686–700.

Sun N, Ma P-C, Yan S, Wen S-Q, Sun C, Du P-X, Cheng H-Z, Deng X-H, Wang C-C, Wei L-H. 2019. Phylogeography of Y-chromosome haplogroup Q1a1a-M120, a paternal lineage connecting populations in Siberia and East Asia. Annals of Human Biology 46: 261–266.

Tao Y, Zhou J, Liang L, Allen E, Zou Y, Huang Z, Li H. 2023. Fine-scale Genetic Structure of Geographically Distinct Patrilineal Lineages Delineates Southward Migration Routes for Han Chinese. Nature Anthropology.

Wang C, Dai J, Qin N, Fan J, Ma H, Chen C, An M, Zhang J, Yan C, Gu Y et al. 2022a. Analyses of rare predisposing variants of lung cancer in 6,004 whole genomes in Chinese. Cancer Cell 40: 1223–1239 e1226.

Wang CC, Li H. 2013. Inferring human history in East Asia from Y chromosomes. Investig Genet 4: 11.

Wang CC, Yeh HY, Popov AN, Zhang HQ, Matsumura H, Sirak K, Cheronet O, Kovalev A, Rohland N, Kim AM et al. 2021a. Genomic insights into the formation of human populations in East Asia. Nature 591: 413–419.

Wang F, Song F, Song M, Li J, Xie M, Hou Y. 2021b. Genetic reconstruction and phylogenetic analysis by 193 Y-SNPs and 27 Y-STRs in a Chinese Yi ethnic group. Electrophoresis 42: 1480–1487.

Wang F, Song F, Song M, Luo H, Hou Y. 2022b. Genetic structure and paternal admixture of the modern Chinese Zhuang population based on 37 Y-STRs and 233 Y-SNPs. Forensic Sci Int Genet 58: 102681.

Wang M, He G, Zou X, Liu J, Ye Z, Ming T, Du W, Wang Z, Hou Y. 2021c. Genetic insights into the paternal admixture history of Chinese Mongolians via high-resolution customized Y-SNP SNaPshot panels. Forensic Sci Int Genet 54: 102565.

Wang MG, He GL, Zou X, Chen PY, Wang Z, Tang RK, Yang XM, Chen J, Yang MQ, Li YX et al. 2022c. Reconstructing the genetic admixture history of Tai-Kadai and Sinitic people: Insights from genome-wide SNP data from South China. Journal of Systematics and Evolution 61: 157–178.

Wang T, Wang W, Xie G, Li Z, Fan X, Yang Q, Wu X, Cao P, Liu Y, Yang R et al. 2021d. Human population history at the crossroads of East and Southeast Asia since 11,000 years ago. Cell 184: 3829–3841 e3821.

Wang Z. 1994. History of nationalities in China. China Social Science Press, Beijing (in Chinese).

Wen B, Li H, Lu D, Song X, Zhang F, He Y, Li F, Gao Y, Mao X, Zhang L et al. 2004a. Genetic evidence supports demic diffusion of Han culture. Nature 431: 302–305.

Wen B, Xie X, Gao S, Li H, Shi H, Song X, Qian T, Xiao C, Jin J, Su B et al. 2004b. Analyses of genetic structure of Tibeto-Burman populations reveals sexbiased admixture in southern Tibeto-Burmans. Am J Hum Genet 74: 856–865.

Xie M, Song F, Li J, Lang M, Luo H, Wang Z, Wu J, Li C, Tian C, Wang W et al. 2019. Genetic substructure and forensic characteristics of Chinese Hui populations using 157 Y-SNPs and 27 Y-STRs. Forensic Sci Int Genet 41: 11–18.

Xu J. 1999. Xueqiu: Han minzu de renleixue fenxi (Snowball: An Anthropological Analysis of the Han Nationality). Shanghai: Shanghai renmin chubanshe.

Xu J. 2012. Understanding the snowball theory of the Han nationality. Critical Han studies: the history, representation, and identity of China’s majority: 113–127.

Xu S, Yin X, Li S, Jin W, Lou H, Yang L, Gong X, Wang H, Shen Y, Pan X et al. 2009. Genomic dissection of population substructure of Han Chinese and its implication in association studies. Am J Hum Genet 85: 762–774.

Xue F, Wang Y, Xu S, Zhang F, Wen B, Wu X, Lu M, Deka R, Qian J, Jin L. 2008. A spatial analysis of genetic structure of human populations in China reveals distinct difference between maternal and paternal lineages. Eur J Hum Genet 16: 705–717.

Xue Y, Zerjal T, Bao W, Zhu S, Shu Q, Xu J, Du R, Fu S, Li P, Hurles ME. 2006. Male demography in East Asia: a north–south contrast in human population expansion times. Genetics 172: 2431–2439.

Yan S, Wang CC, Zheng HX, Wang W, Qin ZD, Wei LH, Wang Y, Pan XD, Fu WQ, He YG et al. 2014. Y chromosomes of 40% Chinese descend from three Neolithic super-grandfathers. PLoS One 9: e105691.

Yang L, Tan S, Yu H, Zheng B, Qiao E, Dong Y, Zan R, Xiao C. 2008. Gene admixture in ethnic populations in upper part of Silk Road revealed by mtDNA polymorphism. Science in China Series C, Life sciences 51: 435–444.

Yang MA, Fan X, Sun B, Chen C, Lang J, Ko YC, Tsang CH, Chiu H, Wang T, Bao Q et al. 2020. Ancient DNA indicates human population shifts and admixture in northern and southern China. Science 369: 282–288.

Yao H, Wang M, Zou X, Li Y, Yang X, Li A, Yeh H-Y, Wang P, Wang Z, Bai J. 2021. New insights into the fine-scale history of western–eastern admixture of the northwestern Chinese population in the Hexi Corridor via genome-wide genetic legacy. Molecular Genetics and Genomics 296: 631–651.

Yao YG, Kong QP, Bandelt HJ, Kivisild T, Zhang YP. 2002. Phylogeographic differentiation of mitochondrial DNA in Han Chinese. Am J Hum Genet 70: 635–651.

Yu C, Lan X, Tao Y, Guo Y, Sun D, Qian P, Zhou Y, Walters Robin G, Li L, Zhu Y et al. 2023a. A high-resolution haplotype-resolved Reference panel constructed from the China Kadoorie Biobank Study. Nucleic Acids Research doi:10.1093/nar/gkad779.

Yu H-X, Ao C, Wang X-P, Zhang X-P, Sun J, Li H, Liu K-J, Wei L-H. 2023b. The impacts of bronze age in the gene pool of Chinese: Insights from phylogeographics of Y-chromosomal haplogroup N1a2a-F1101. Frontiers in Genetics 14: 1139722.

Yu H, Spyrou MA, Karapetian M, Shnaider S, Radzeviciute R, Nagele K, Neumann GU, Penske S, Zech J, Lucas M et al. 2020. Paleolithic to Bronze Age Siberians Reveal Connections with First Americans and across Eurasia. Cell 181: 1232–1245 e1220.

Zhang F, Ning C, Scott A, Fu Q, Bjorn R, Li W, Wei D, Wang W, Fan L, Abuduresule I et al. 2021a. The genomic origins of the Bronze Age Tarim Basin mummies. Nature 599: 256–261.

Zhang M, Yan S, Pan W, Jin L. 2019. Phylogenetic evidence for Sino-Tibetan origin in northern China in the Late Neolithic. Nature 569: 112–115.

Zhang P, Luo H, Li Y, Wang Y, Wang J, Zheng Y, Niu Y, Shi Y, Zhou H, Song T et al. 2021b. NyuWa Genome resource: A deep whole-genome sequencing-based variation profile and reference panel for the Chinese population. Cell Rep 37: 110017.

Zhong H, Shi H, Qi XB, Duan ZY, Tan PP, Jin L, Su B, Ma RZ. 2011. Extended Y chromosome investigation suggests postglacial migrations of modern humans into East Asia via the northern route. Mol Biol Evol 28: 717–727.

Zhou Y, Zhan X, Jin J, Zhou L, Bergman J, Li X, Rousselle MMC, Belles MR, Zhao L, Fang M et al. 2023. Eighty million years of rapid evolution of the primate Y chromosome. Nat Ecol Evol 7: 1114–1130.

